# Multiomic Profiling of 85 iPSC Lines from Familial Dementia Reveals Cellular Diversity and Regulators of Organoid Development

**DOI:** 10.64898/2026.04.29.721747

**Authors:** Le Qi, Taylor Bertucci, Susan Borden, Brigitte L. Arduini, Steven Lotz, Kathryn Bowles, Alison Goate, Celeste M. Karch, Sally Temple, Daniel H Geschwind

## Abstract

Patient-derived induced pluripotent stem cells (iPSCs) are used for disease modeling and therapeutic development, yet their systematic molecular and genetic characterization is uncommon. Here, we performed whole genome sequencing (WGS), bulk/single-cell RNA-sequencing (RNA-seq) and DNA methylation analysis on 85 iPSC lines harboring *MAPT* mutations and corresponding isogenic controls. Single-cell RNA-seq revealed reproducible inter- and intra-line heterogeneity related to quality/pluripotency and nominated surface markers to quantify iPSC subclusters. WGS detected unintended editing events and structural variants, including the 20q21.31 duplication, missed by standard assays. We identified pathways correlated with neural organoid formation efficiency and with genome-editing and clonal-selection effects, underscoring the need to use unedited lines as isogenic controls. This comprehensive dataset improves the utility of the MAPT iPSC collection and provides proof of principle supporting in-depth genomic characterization to improve iPSC utility in biological research.

## INTRODUCTION

Patient-derived induced pluripotent stem cells (iPSCs) offer unique opportunities to study how genetic risk factors cause human disease. Scaled differentiation into disease-relevant and often inaccessible cell types supports high-throughput functional assays that recapitulate many *in vivo* pathophysiological features^1–13^. However, considerable heterogeneity has been observed across iPSC lines, including differences in pluripotency extent, differentiation potential, and molecular characteristics such as genomic and transcriptomic landscape^14–16^. This heterogeneity poses a significant challenge for iPSC-derived models, increasing experimental variability and obscuring disease-relevant biological differences. Yet comprehensive molecular and genetic characterization of lines is seldom performed, and few large-scale analyses of heterogeneity and potential confounding mutations introduced during iPSC editing or culturing have been conducted. Moreover, because patient mutations are rare, individual labs or institutions typically lack sufficiently broad collections to compare donors with the same mutation, to compare across different mutations, or to enable independent validation. To address this gap, the Tau Consortium Stem Cell Group generated more than 100 iPSC lines with dominantly inherited *MAPT* mutations that cause a range of frontotemporal dementia syndromes (FTDs) and corresponding genome-engineered isogenic controls^17–20^ for the study of human neurodegenerative disease mechanisms. These lines have been distributed to over 70 laboratories and have supported numerous publications^21–37^.

Complicating the use of iPSC lines to understand disease mechanisms is the emergence of *de novo* mutations, especially structural variations, which often occur during reprogramming, passaging, and genome editing of iPSCs^38–43^. Although this problem can be partially mitigated by using independent pairs of isogenic mutants and wild-type controls, a more complete picture of the iPSC genome would enable more effective selection of iPSC lines for study and more careful interpretation of experimental results. We reasoned that comprehensive genomic profiling would offer an essential resource not only for those using these lines but also provide generalizable insights into iPSC biology and avenues for improving directed differentiation. Beyond neurodegeneration, we considered that this iPSC line resource and its comprehensive molecular and genomic characterization (e.g., WGS, transcriptomics, methylation, long-read sequencing), would also deliver critical value for diverse fields by revealing off-target edits, structural variants, clonal and cell-state heterogeneity, and biomarkers, improving reproducibility, enabling cross-disease comparisons, and ultimately strengthening target validation and translational pipelines.

To systematically characterize the genomic, epigenomic, transcriptomic profiles and cellular heterogeneity of these iPSC lines, we performed whole genome sequencing (WGS), long-read DNA sequencing of the 17q21.31 *MAPT* locus, bulk and single-cell RNA sequencing (RNA-seq), and measured bulk DNA methylation via microarray.

Combining these multi-modal datasets allowed us to generate a comprehensive molecular profile of these widely used iPSC lines. We correlated these quantitative measurements performed at the iPSC stage with available cellular phenotypes to gain insight into the processes that regulate the efficiency of iPSC differentiation into neural organoids and the signaling pathways altered during CRISPR editing.

## RESULTS

### Description of the iPSC cohort and molecular profiling

Tau deposition, or tauopathy is a central feature of multiple neurodegenerative disorders. These include primary tauopathies, where tau deposition is the primary driver, and secondary tauopathies, such as Alzheimer’s disease, where pathogenic tau deposition is considered a consequence of other causal factors. We profiled 85 human iPSC lines encompassing 9 different *MAPT* mutations associated with primary tauopathies, and isogenic controls derived from 25 unique donors of European ancestry (Figure 1A). Twenty donors have *MAPT* mutations, two have idiopathic progressive supranuclear palsy (PSP) and three donors represent healthy controls. Multiple parental clones were included for 14 donors. Isogenic clones that were either edited (Figure S1A) or unedited (Figure S1A) after parallel CRISPR editing and clonal expansion were also included for 21 donors to control for the effects of the editing procedure itself. In parallel, several *MAPT* mutations were introduced into two control lines to support isogenic comparisons (Figure S1B).

**Figure 1:**
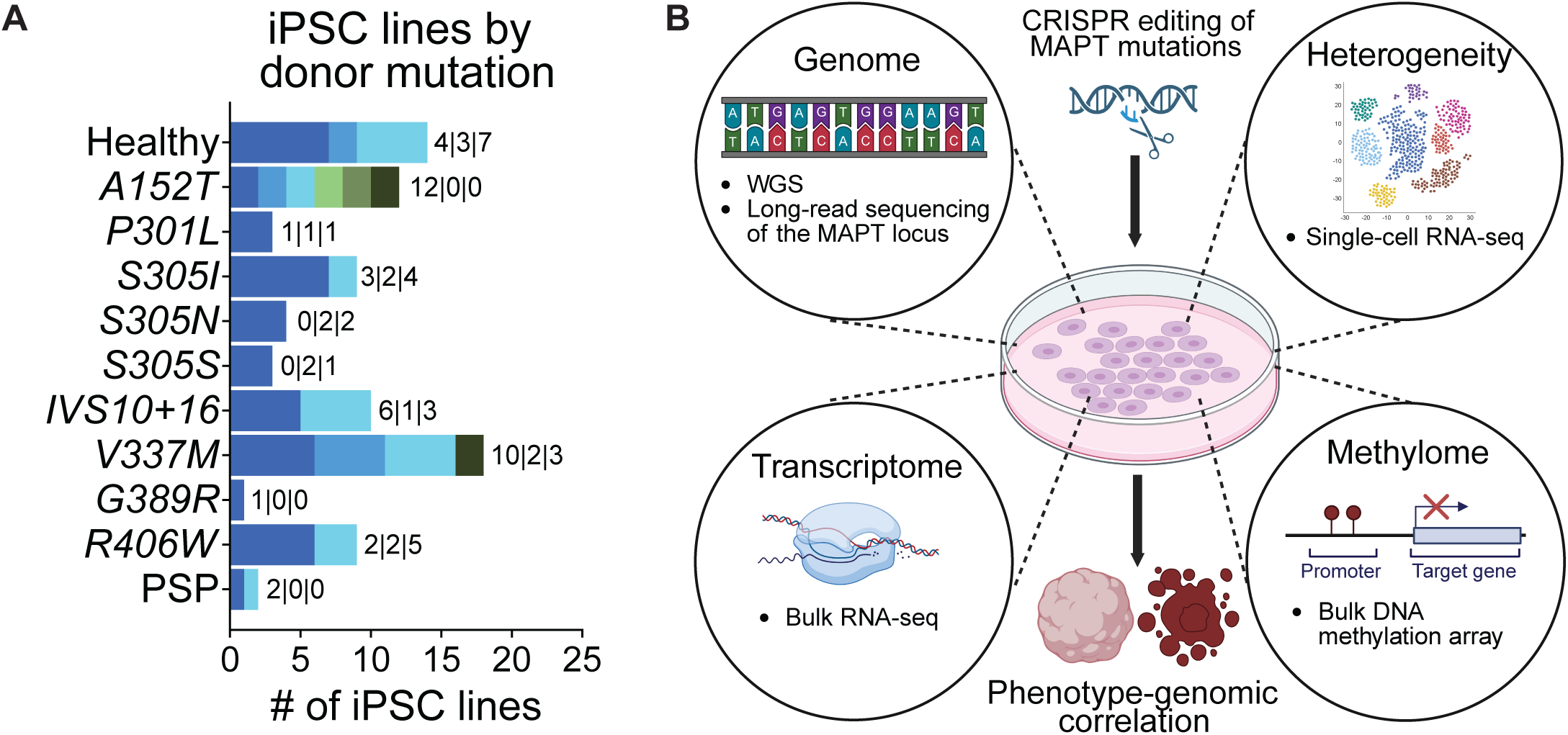
Overview of the iPSC cohort and molecular profiling performed (A) iPSC lines in this study grouped by *MAPT* mutations present in their donors. The numbers of unique lines and donors (in bracket) are specified for each mutation. (B) Molecular profiling performed on this iPSC cohort, including WGS, long-read DNA sequencing of the MAPT locus, single-cell RNA-seq, bulk RNA-seq and bulk DNA methylation array. These profiling results were associated with iPSC phenotypes, such as the efficiency of neural organoid formation and CRISPR editing status. See also Figure S1 and Table S1.

We cultured these 85 iPSC lines in 9 separate batches, with isogenic lines grouped together and one control line F11350-1 included in every batch for comparison and normalization. We selected low-passage-number clones, with mean passage number of 15.9 ± 6.4 for parental lines, 34.7 ± 12.7 for unedited lines, and 32.5 ± 10.5 for edited lines (Figure S1C). We used lot-matched culture reagents to minimize culture batch effects. All samples were cultured in mTeSR medium supplemented with FGF2-DISCs, with 13 lines cultured in standard daily fed mTeSR medium without FGF2-DISCs in parallel^44^. Extensive quality control was performed on these lines, including karyotyping, mycoplasma testing, removal of samples with spontaneous differentiation, and examination of pluripotency/differentiation markers (Figures S1D and S1E). The detailed metadata for this collection of iPSCs are provided in Table S1. We performed 30x WGS, long-read DNA sequencing of the *MAPT* locus on the PacBio platform, bulk and single-cell RNA-seq, and measured bulk DNA methylation via microarray (Figure 1B; Methods). We used these data to understand line quality and heterogeneity and gain further biological insight into the mechanisms underlying their behavior.

### Cellular and transcriptional heterogeneity within iPSC lines

Inter-clonal heterogeneity of iPSCs has been well documented, mostly driven by donor genetic background^45–51^. However, intra-clonal heterogeneity at the undifferentiated iPSC level has been largely understudied. Nguyen *et al*. performed scRNA-seq on a single WTC11-CRISPRi iPSC line and identified 4 clusters ranging from a core pluripotent state to late primed state^52^. However, it is difficult to draw generalizable conclusions based on a single iPSC line. Here, we leveraged this large collection of iPSC lines to identify consistent patterns of intra-clonal heterogeneity using a robust analysis pipeline (Figure S2A). Using single cell RNA-seq (scRNA-seq), we profiled 2713 ± 1314 cells per line after doublet removal and stringent QC filtering (Methods), identifying 4882 ± 1777 genes per cell (Figures S2B-S2D). We bootstrapped 80% of the cells 20 times to choose stable clustering parameters (Figure S2A; Methods) This resulted in 4 stable clusters in most lines, with 3 of which present in all lines (Figures 2A and 2B). To annotate these clusters, we mapped our data to the Human Primary Cell Atlas dataset^53^. As expected, the vast majority of the cells matched with either iPSCs or embryonic stem cells (ESCs), while the smallest cluster was mapped to ESC-derived mesodermal and neural lineages, suggesting an early differentiation-poised state (hereafter referred to as “poised iPSCs”, Figure S2E).

**Figure 2:**
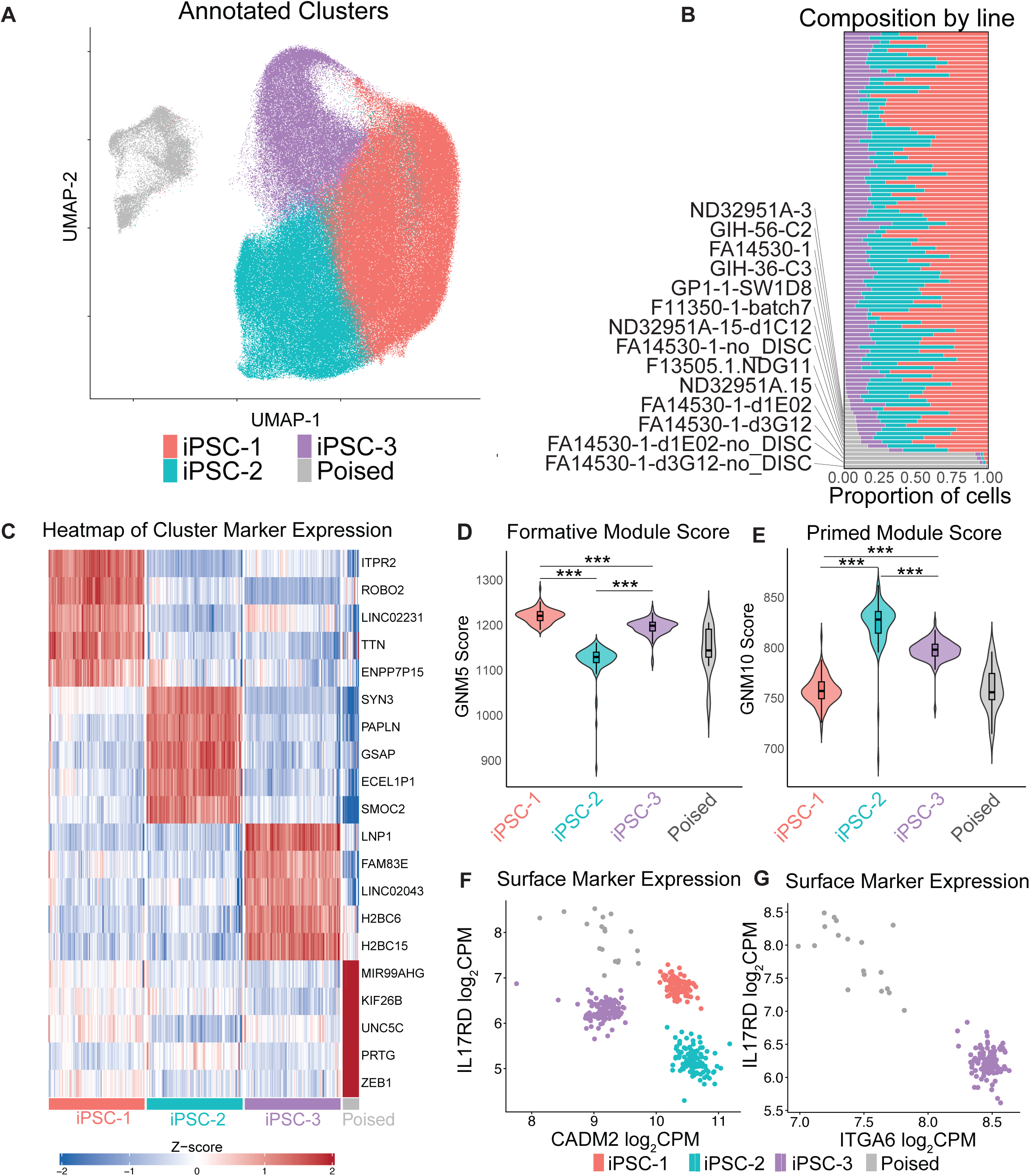
Cellular and transcriptional heterogeneity within iPSC lines (A) Uniform Manifold Approximation and Projection (UMAP) representation of 4 Leiden clusters. The colors for each cluster are shared throughout this figure. (B) The proportion of each Leiden cluster for each iPSC line is ordered by the proportion of the poised cluster. iPSC lines with >5% poised iPSC cluster are labeled. (C) Heatmap of top 5 marker genes significantly up-regulated in each Leiden cluster versus all other clusters (D and E) Violin plots of gene network module (GNM) scores for formative module (GNM5) and poised module (GNM10) as described in Arthur *et al.* for each iPSC clusters (Methods).^54^ Paired t-test with FDR corrections were performed only among iPSC-1, iPSC-2 and iPSC-3 clusters. (F and G) Scatter plots of gene expression values (log_2_CPM) of identified surface marker genes distinguishing Leiden clusters. Cells with CADM2 log_2_CPM < 10 (D) are isolated and plotted in E. See also Figure S2 and Table S2.

We next performed analysis of differential gene expression across clusters in a pseudo-bulk fashion (Figure S2A, Methods). We identified 1499, 1983, 5846, and 2029 genes significantly up-regulated in iPSC-1, iPSC-2, iPSC-3, and poised iPSC clusters, respectively (Figure 2C and Table S2A). Pluripotency genes POU5F1 (OCT4), NANOG and SOX2 were expressed in all clusters, even though the expression levels varied (Figure S2F). Consistent with reference mapping results (Figure S2E), poised iPSC clusters showed significantly lower and more variable expression of these pluripotency genes (Figure S2F). To understand the pluripotency states of iPSC-1, iPSC-2 and iPSC-3 clusters, we calculated the module scores associated with formative pluripotency state (a state between naïve and primed pluripotency, as measured by gene network module (GNM) 5 score as described in Arthur *et al.* (2024)^54,55^, Figure 2D) and primed pluripotency state (GNM 10, Figure 2E). iPSC-1 most resembled the formative state, followed by iPSC-3, whereas iPSC-2 showed the highest primed pluripotency signature (Figures 2D and 2E). Corroborating these results, genes significantly up-regulated in iPSC-1 compared to the rest of the clusters were enriched in cell cycle pathways, consistent with the faster cell cycle rate associated with the formative state (Figure S2G top left and Table S2B)^55^. Neural differentiation and synaptic function pathways were enriched in iPSC-2, consistent with their more primed state (Figures 2E, S2G top right, and Table S2B). Finally, the poised iPSC cluster showed enrichment of several lineage differentiation pathways, also consistent with the reference mapping results (Figure S2G bottom right, S2E and Table S2B).

As shown in Figure 2B, the proportions of the iPSC-1, iPSC-2 and iPSC3 clusters were relatively stable across lines. However, we observed that the proportion of the poised iPSC cluster (gray) varied significantly across lines, being absent in most, but more abundant in a small subset of lines (Figure 2B). To identify the variables contributing to the compositional differences in an unbiased manner, we performed variance partitioning^56^. Donor explained the most compositional variance, especially in the poised iPSC cluster (Figure S2H). In particular, lines derived from one donor (FA14530), which had very poor organoid formation capacity (Table S1), stood out for having a disproportionally high percentage of poised iPSC cluster (Figure 2B; Methods). As we have previously shown, adding FGF2-DISCs for sustained release improves the quality of iPSCs^57^, which correlated with a significant reduction of the poised iPSC percentage (Figure S2I). The proportion of the poised iPSC cluster did not significantly correlate with either the passage number (Figure S2M), or the percentage of differentiation marker SSEA1^+^ cells (Figure S2N)^58^.

These results suggest that the proportion of the poised iPSC cluster reflects an orthogonal aspect of iPSC quality, which is not captured by existing QC measurements, but easily visible in single cell transcriptome data. As a result, we set out to identify cell surface markers to distinguish these iPSC clusters. As an example, the combination of *CADM2*, *IL17RD* and *ITGA6* expression can effectively separate all 4 clusters (Figures 2F and 2G). Additional marker combinations are included in Table S2C. Thus, scRNA-seq revealed consistent intra-clonal heterogeneity within seemingly homogeneous iPSC lines, the associated gene expression differences, their relationship to iPSC quality, as measured by neural organoid formation and identified surface markers that are candidates for their quantification as part of comprehensive QC.

### Significant de novo putative protein-altering variants (pPAVs) are infrequent, despite occasional large CRISPR on-target deletions

Next, we sought to examine if any significant variants were present in this cohort either inherited from the donor, or spontaneously arising during iPSC manipulation. We performed WGS at 30x coverage (Figure S3A) and used standard pipelines to jointly call short variants (SNPs and INDELs < 50 bp) and structural variants (SV > 50 bp) in all the lines (Figure S2B; Methods). As expected, all lines derived from the same donors clustered together (Figure 3A). However, we discovered that two of the donors, 75-11 and MHF-110, had an identical genetic background and the same *MAPT* mutation (Figure 3A and Table S1), indicating either a sample mix-up or duplicated donation. Additionally, we performed deep long-read DNA sequencing targeting the *MAPT* locus, given its complex genetic structure (Figures S3C and S3D)^59,60^. We compared the *MAPT* variants called from sequencing results to their records and identified 5 lines with inconsistent *MAPT* genotypes (Figure 3B). In particular, GP1-1-WAC11 and 75-11-IW1B9 share the same 10bp out-of-frame deletion at the 5’ end of the original *S305* mutation, potentially resulting from the same CRISPR editing strategy. FA14530-1-d1G12, which was thought to be a corrected isogenic WT control derived from its parental *R406W* line, had a 156bp in-frame deletion that removed the region containing the original *R406W* mutation (Figure 3C), which was missed by Sanger sequencing of the locus. These results highlight the importance of more extensive genetic characterization after CRISPR editing and routine confirmation of genotype of interest^61^. Overall, this initial characterization by WGS led to re-annotation of 5 lines (Figure 3B).

**Figure 3:**
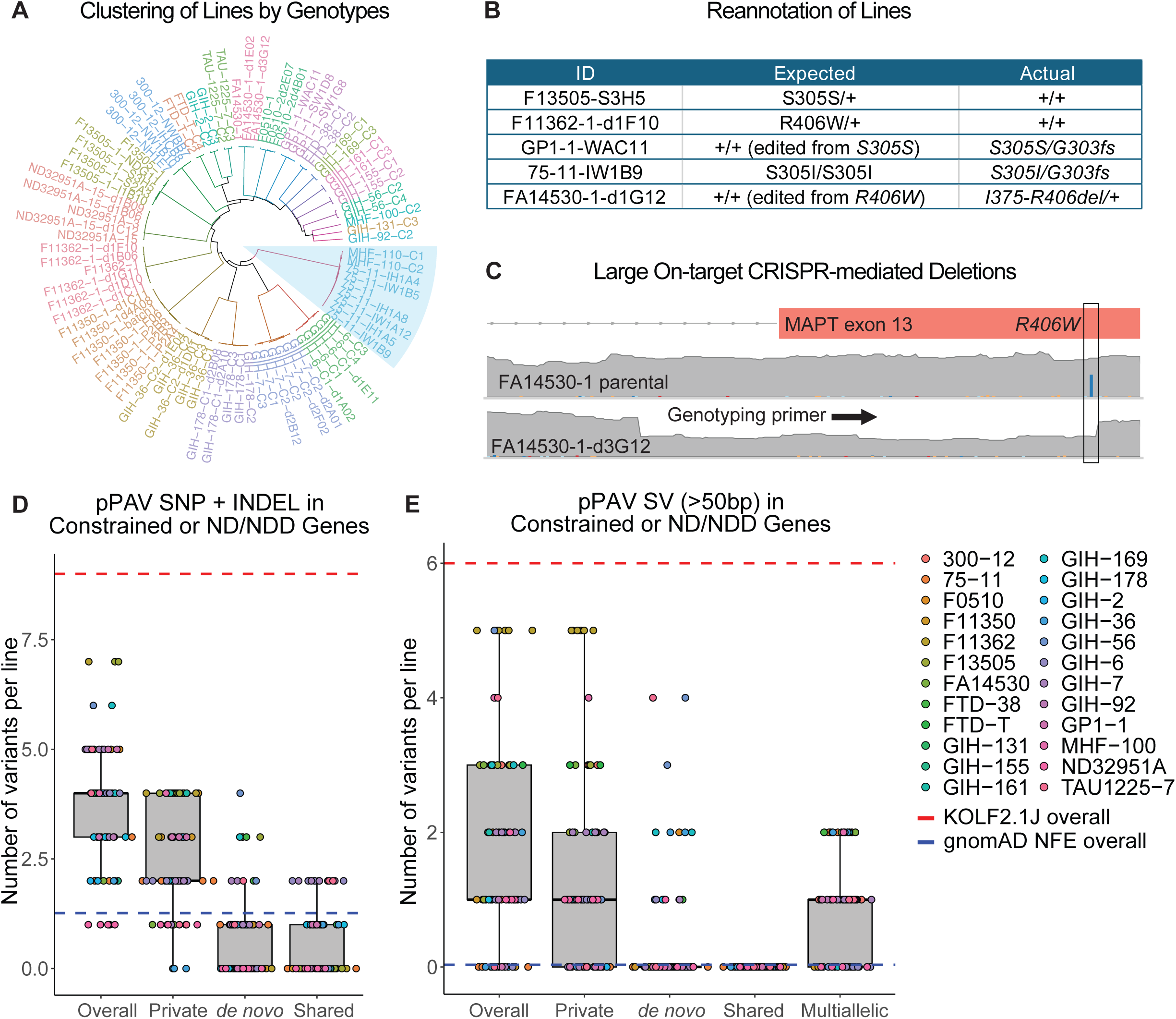
Significant de novo protein-altering variants (pPAVs) are infrequent, despite occasional large CRISPR on-target deletions (A) Dendrogram of hierarchical clustering using SNPs and short INDELs (< 50bp) called in each iPSC line. The area with light blue shade indicates identical genetic background shared by lines derived from donors MHF-110 and 75-11. (B) List of lines with *MAPT* genotypes reannotated according to sequencing data. (C) coverage plot depicting a 156bp deletion surrounding *R406W* mutation in FA14530-1-d3G12 line. The black box indicates the location of the *R406W* mutation in the parental line. The arrow indicates the primer used for genotyping, which was missing after the deletion. (D and E) Box plots of putative protein-altering variants (pPAVs) in constrained or neurodegenerative/neurodevelopmental disorder (ND/NDD) genes in each line called as SNPs and short INDELs (< 50bp) (D) and structural variants (SVs, > 50bp) (E). Variants were further grouped by their frequency among iPSC lines, namely private (present in > 50% of the lines derived from a single donor), *de novo* (< 50% of the lines derived from a single donor), shared (> 1 donor) and multiallelic (multiallelic copy number variations). Known *MAPT* variants either derived from donors or introduced by CRISPR-editing were excluded. Red and blue dashed lines indicate total numbers of corresponding variants in KOLF2.1J iPSC line^62^ and average number of variants in non-Finish European population in gnomAD v4.1. See also Figure S3 and Table S3.

To expand the analysis of functionally relevant variants to a genome-wide scale, we quantified the number of different forms of variants by frequency and impact, including overall variants (Figures S3E and S3F), rare variants (minor allele frequency (MAF) < 0.01% in any gnomAD populations, Figures S3G and S3H), rare, putative protein-altering variants (pPAVs, Figures S3I and S3J), as well as rare pPAVs in constrained (LOEUF < 0.6, gnomAD v4.1; Methods) or neurodegenerative/neurodevelopmental disorder (ND/NDD) risk genes (Figures 3D and 3E, Table S3; Methods). We categorized these variants into 4 groups, those present in: a) >50% of the lines derived from the same donor (private), b) <50% of the lines from the same donor (*de novo*), c) >1 donor (shared), or d) multiallelic CNVs. We benchmarked the number of pPAVs in constrained or ND/NDD genes in our cohort with published WGS data from the KOLF2.1J iPSC line selected for the iPSC Neurodegenerative Disease Initiative (iNDI) from the NIH’s Center for Alzheimer’s and Related Dementias (CARD)^62^, as well as the non-Finnish European (NFE) population in gnomAD v4.1 (red and blue dashed line in Figures 3D and 3E). We observed slightly more overall variants per line as compared to gnomAD NFE population averages (blue dashed line, Figures 3D and 3E). This difference likely reflects a combination of ascertainment bias toward tauopathy donors and the accumulation of *de novo* mutations during iPSC derivation and culture. When compared with the KOLF2.1J WGS data processed in the same manner as ours, this cohort contained approximately 65% fewer variants than the KOLF2.1J line^69^. The majority of these variants in our cohort were private, which suggests that they are most likely germline mutations, but we do not have access to donor DNA WGS data to confirm this. We did not detect any cancer-associated point mutations previously identified as recurrent in human iPSC cultures^41,42^, the only exception being the 20q11.21 duplication in approximately 20% of the lines (hereafter referred to as 20q DUP, Table S3), as discussed below.

### The recurrent 20q11.21 duplication up-regulates ID1 and impairs neural organoid formation

Although all iPSC lines passed the karyotyping QC step, the 1 Mb duplication in this region is smaller than the typical resolution of karyotyping (>3-5 Mb). As expected, we observed that the 20q11.21 copy number is significantly correlated with passage number (Figure 4A; Spearman ρ=0.54, p=5.24e−08), due to the strong selective advantage it confers via *BCL2L1* up-regulation^63,64^. To study the functional consequences of 20q DUP in iPSCs, we performed bulk RNA-seq and DNA methylation array (Illumina 850K Infinium MethylationEPIC) on this cohort, controlling for passage number (Figure 4A; Methods). Similar to previous reports^65^, donor and batch contributed the most to variance in both gene expression and DNA methylation, given that isogenic lines were cultured in the same batches (Figures S4A and S4B). The methylation age^66^ of the lines also correlated strongly with their passage numbers, as previously reported (Figure S4C). Lines with 20q DUP had significant up-regulation of genes located in this locus, including *HM13*, *ID1*, *BCL2L1*, *TPX2 and PDRG1*, consistent with previous studies^63,67,68^ (Figure 4B). To examine whether the 20q DUP impairs neural differentiation as previously described^67,68^, we tested the efficiency to form well-patterned neural organoids in a subset of lines^57^. Indeed, half of the lines that had been identified as yielding poorly patterned neural organoid (defined as the absence of FOXG1 staining at day 20 post differentiation, referred to as poor lines hereafter) harbored the 20q DUP, while all efficient lines did not (Figure 4C and Table S1). Interestingly, the FA14530 lines, which exhibited a high proportion of poised iPSC clusters (Figure 2B) and poor capacity to form neural organoids (Table S1), did not harbor the 20q DUP. This suggests that multiple independent factors affect organoid formation capacity.

**Figure 4:**
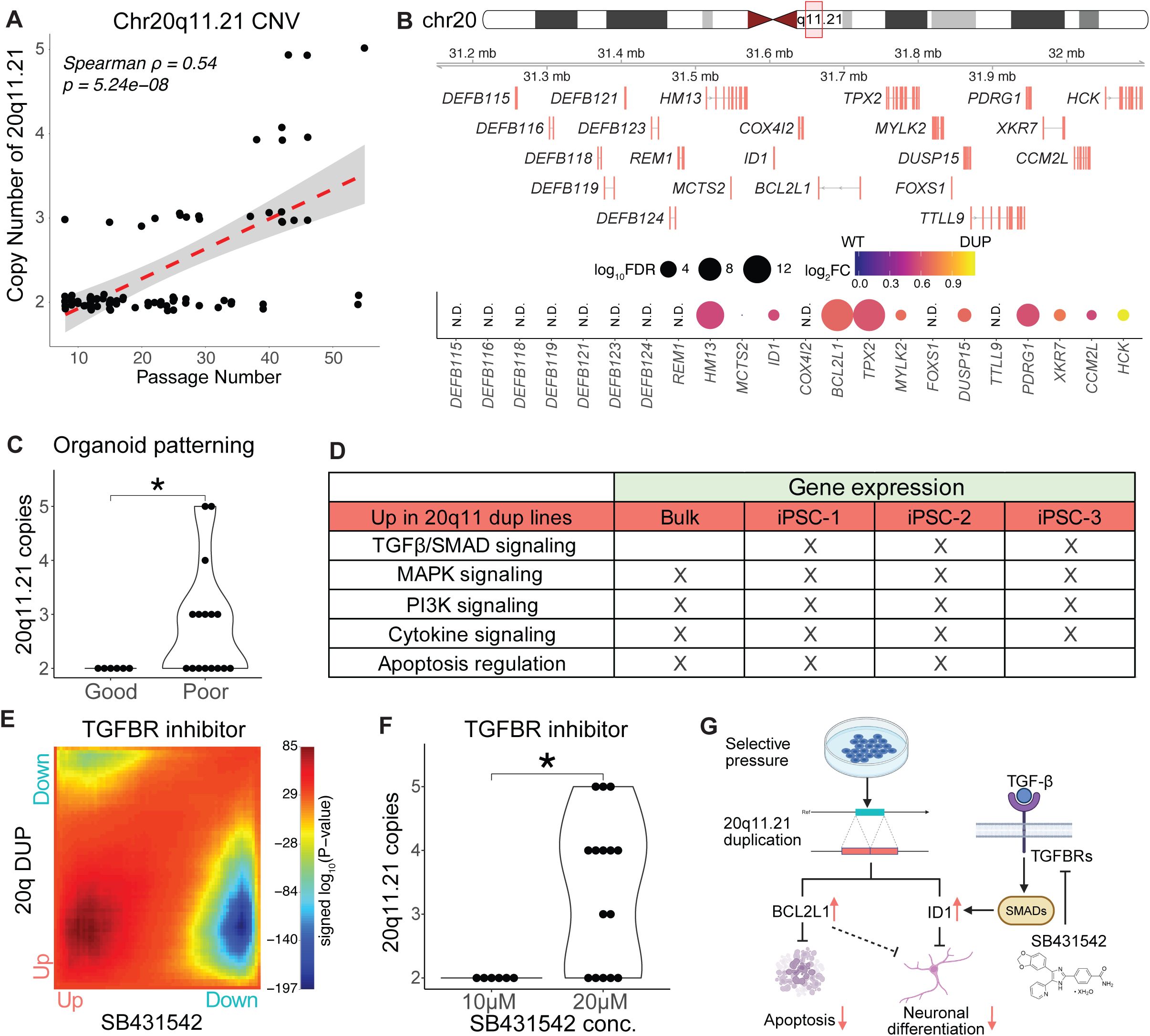
Recurrent 20q11.21 duplication up-regulates ID1 and impairs neural organoid formation (A) Correlation between the copy number of 20q11.21 and passage number for each iPSC line. Data points were jiggled to minimize overlap. (B) Dot plot representing local transcriptional changes associated with 20q DUP in conjunction with transcript annotations of this region. (C) Number of 20q11.21 copy numbers in lines determined to have efficient and inefficient neural organoid formation. (D) Pathways consistently changed in GSEA results comparing lines with and without 20q DUP across RNA-seq results (bulk and iPSC clusters). (E) Heatmap of rank ranked hypergeometric overlap (RRHO) analysis^85^ comparing transcriptomic changes associated with 20q DUP and 24h SB431542 treatment in H9 human embryonic stem cells^82^. Positive p-values indicates positive correlation and vice versa. (F) Number of 20q11.21 copy numbers in lines determined to require double SB431542 concentration during neural organoid formation. (G) Schematics of our working model depicting the role of ID1 in neuronal differentiation and its interaction with TGF-β pathway inhibition during organoid formation. Wilcoxon test was used for panel E and H. See also Figure S4 and Table S4.

The upregulation of *BCL2L1,* which is located in the 20q11.21 region, has been shown to contribute to impaired neural differentiation in iPSC models.^68^ But, the potential role of other genes in this region has not been underexplored. In particular, ID1 plays an essential role in maintaining pluripotency and preventing neural differentiation^69–72^, making it a plausible contributor. To identify factors contributing to impaired organoid formation, we performed gene set enrichment analysis (GSEA) comparing lines with and without 20q DUP in bulk methylation, bulk RNA-seq and scRNA-seq data (Table S4). Minimal differences were observed in terms of cluster composition or DNA methylation (Figures S4D and S4E). 20q DUP led to up-regulation of the “apoptotic regulation pathway” likely due to *BCL2L1* amplification. MAPK and phosphatidylinositide-3 kinases (PI3K) signaling also increased, potentially contributing to their reduced dependence on FGF signaling^73^ (Figure 4D). Intriguingly, TGF-β/SMAD signaling, which plays an essential role in iPSC self-renewal^74–77^, neural differentiation^78,79^ and induces transcription of *ID1* in human iPSCs^80,81^, increased significantly in conjunction with the 20q DUP (Figure 4D). We independently validated that expression of *ID1*, along with the canonical SMAD2/3 target gene *NANOG*, were both significantly reduced in human iPSCs after 24h SB431542 (TGF-β receptor inhibitor) treatment^82^, but significantly increased with *SMAD2* or *SMAD3* over-expression^83^ (Figure S4F). Notably, ID1 and BCL2L1 up-regulation have been shown to synergistically promote iPSC competitive advantage and fate bias^84^. We hypothesized that more copies of *ID1* make iPSCs more sensitive to TGF-β signaling; therefore lines with 20q DUP require a higher concentration of SB431542, which is part of the standard dual inhibition protocol for neuronal differentiation^78^. Consistent with this, transcriptomic changes after 24h of SB431542 treatment^82^ had a strong inverse correlation with 20q DUP in our cohort (rank-rank hypergeometric overlap (RRHO) analysis^85^, Figure 4E; Methods). We also tested whether doubling the SB431542 concentration would improve organoid formation in a subset of the lines with or without 20q DUP. Indeed, organoid formation improved by doubling the SB431542 concentration in all lines with 20q DUP, while having no significant effect in WT lines (Figures 4F)^57^. This suggests that *ID1* contributes to impaired neural differentiation associated with 20q DUP in addition to *BCL2L1,* which is supported by our observation that increasing SB431542 concentration mitigated the effects of the duplication (Figure 4G)^57^.

### Identification of molecular targets to improve neural organoid formation efficiency

To understand pathways regulating neural organoid formation efficiency of iPSCs in a broader context, we directly compared gene expression and DNA methylation differences between lines with well- and poorly-patterned neural differentiation capacity, while controlling for 20q DUP status (Methods). The gene expression differences were further analyzed with GSEA and compared with iPSC perturb-seq^86^ to identify pathways regulating neural organoid formation (Figure 5A). Top up-regulated pathways in good lines included synaptic transmission, ion transport and GPCR signaling pathways (Figures 5B, S4G and S4H). Integrin signaling and collagen metabolism genes, which are closely linked to neural differentiation^87,88^, were also up-regulated in good lines. In contrast, genes related to mitochondrial function, ribosomal biogenesis and cell cycle/DNA replication^89–91^ were down-regulated in good lines (Figure 5B). Corresponding changes in promoter methylation were also observed in genes related to the above-mentioned pathways, suggesting that DNA methylation might be partially driving these transcriptional changes (Figure 5B). For example, genes with increased promoter methylation were enriched in aerobic respiration and cell cycle pathways, which might drive down-regulation of expression of these genes in good lines (Figure 5B). Conversely, genes with decreased promoter methylation were enriched in synaptic transmission, cell-cell adhesion and ion transport pathways, potentially leading to increased expression of these genes in good lines (Figure 5B). Interestingly, compared to control treatment, SB431542 treatment in iPSCs^82^ also led to reduced expression of genes related to mitochondrial metabolism, protein translation, and DNA replication pathways (Figure S4I), consistent with its effects on lines without 20q DUP (Figure 4F).

**Figure 5:**
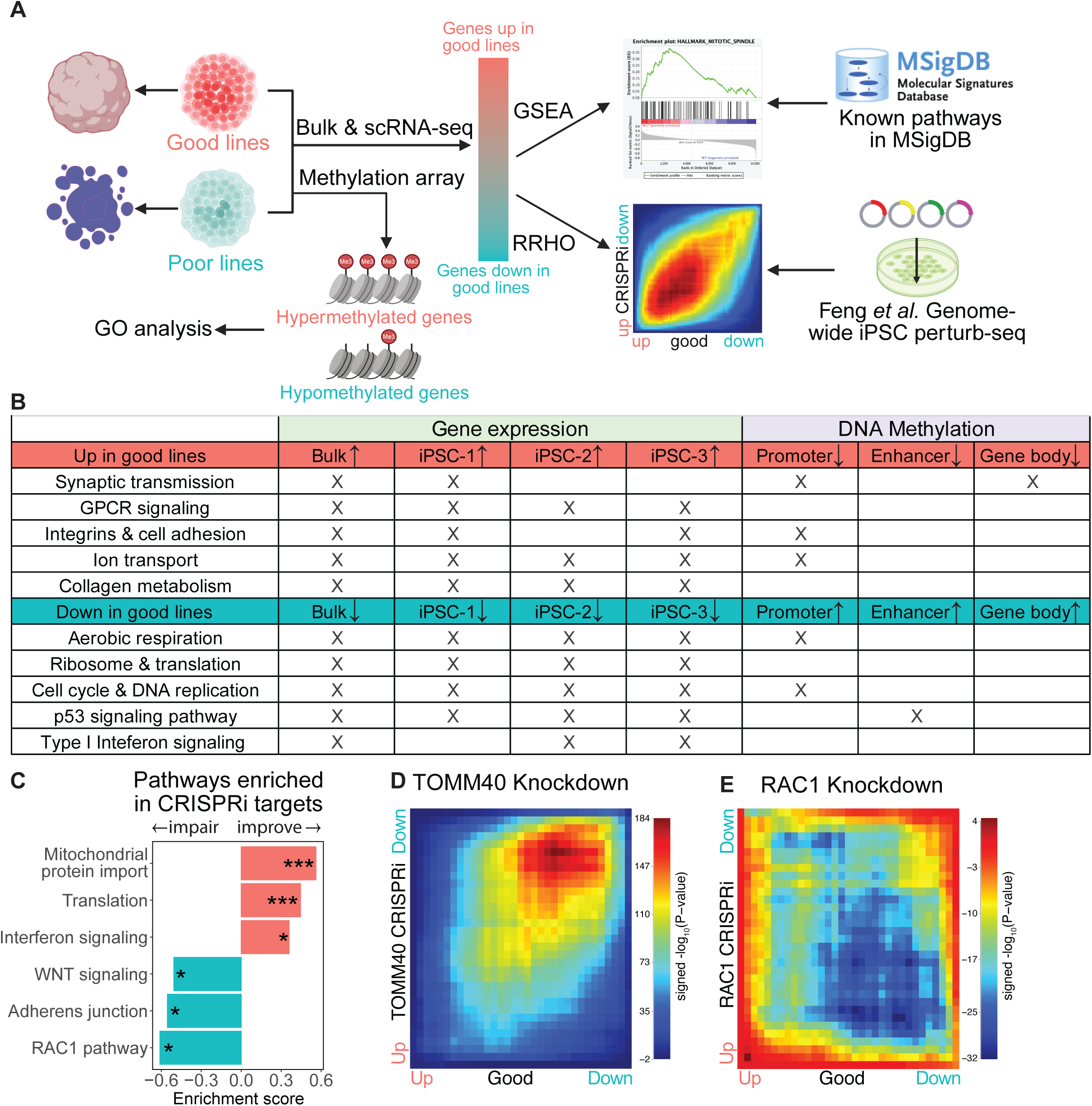
Identification of molecular targets to improve neural organoid formation efficiency (A) Schematics of the analysis performed to understand molecular pathways regulating neural organoid formation. Bulk and single-cell RNA-seq results were compared between efficient and inefficient lines with GSEA performed. Transcriptional changes from bulk RNA-seq were also compared to genome-wide perturb-seq results in human iPSCs described in Feng *et al.*^86^ using RRHO analysis. DNA methylation differences were analyzed and gene ontology (GO) associated with differentially methylated genes were identified. (B) Pathways consistently changed in GSEA results across RNA-seq results (bulk and iPSC clusters) comparing efficient and inefficient lines, which also match GO terms associated with DNA methylation changes. (C) Selected pathways significantly changed in GSEA results using pre-ranked list of genes in Feng *et al.*^86^, the CRISPRi-mediated knockdown of which induced transcriptional changes positively (improve) or negatively (impair) correlates with transcriptional changes associated with efficient lines. (D) RRHO analysis comparing transcriptomic changes associated with efficient lines and TOMM40 knockdown human iPSCs in Feng *et al.*^86^. Positive p-values indicates positive correlation and vice versa. (E) RRHO analysis comparing transcriptomic changes associated with efficient lines and RAC1 knockdown human iPSCs in Feng *et al.*^86^. Positive p-values indicates positive correlation and vice versa. See also Figure S4 and Table S4.

To validate these findings using orthogonal data and gain further insights into molecular pathways regulating organoid formation efficiency, we correlated gene expression changes between good and poor lines against genome-wide perturb-seq results in iPSCs^86^ using RRHO analysis (Figure 5A, Methods). We performed GSEA by ranking CRISPRi targets from the most positive to most negative correlation based on their effects on iPSC transcriptome. Consistent with our observations identifying pathways differentially regulated pathways in good versus poor lines (Figure 5B), knocking-down genes in “mitochondrial protein import” and “protein translation” induced gene expression changes similar to good lines (Figures 5C and 5D). By comparison, knocking-down genes in the “canonical WNT signaling” and “adherens junction pathways” pushed the transcriptome towards patterns observed in poor lines (Figure 5C and 5E). Supporting this observation, existing data shows that increasing WNT pathway inhibitor concentration can lead to poor organoid health^57^. Collectively, the transcriptomic analysis suggests that poor lines are more resistant to neural differentiation, at least partially due to increased mitochondrial and protein synthesis pathways.

### CRISPR editing & clonal selection alters p53 signaling, interferon response and differentiation

We next leveraged the large number of isogenic lines with or without CRISPR editing in this cohort to examine the effects of CRISPR-mediated genome editing and subsequent clonal selection on transcription and DNA methylation in iPSCs. iPSC lines were divided into three groups: 48 parental lines, which were directly derived from the donors and did not undergo genome editing; 25 edited lines, which went through CRISPR engineering process to either correct the original *MAPT* mutations or to introduce new *MAPT* mutations in healthy controls; and 15 unedited lines, which underwent the CRISPR editing processes in parallel with the edited lines, but remained unedited at the target loci and served as controls (Figure 6A). We included passage number and 20q DUP status in our model given that parental lines have lower passage number on average (Figure S1C). Parental lines were largely separated from the other two groups, while edited and unedited lines were indistinguishable from each other (Figure 6B). Consistent with this result, no significant DEGs were found between edited and unedited lines (Figure 6C), which suggested that the editing process had similar impacts whether or not it was successful. Indeed, comparisons of parental lines with combined converted and converted lines (hereafter referred to as clonal lines), showed many significant DEGs and a slightly higher proportion of iPSC-2 clusters, but few DNA methylation differences after controlling for differences in passage numbers and 20q DUP status (Figures 6D, S4J and S4K). Notably, clonal lines showed decreased “p53/DNA damage signaling” and the “interferon response pathway”, which are cellular responses to CRISPR-mediated DNA double-stranded breaks^92,93^ and introduction of exogenous nucleic acids^94,95^, respectively (Figure 6E). Collectively, these results highlight that isogenic controls that have undergone the same genome editing procedures as those that are edited have shared changes due to the experimental procedure and are distinct from parental lines, even when passage numbers are matched. These findings highlight the importance of using controls that have undergone genome editing and clonal selection, rather than parental lines, for comparisons between mutant and non-mutant lines.

**Figure 6:**
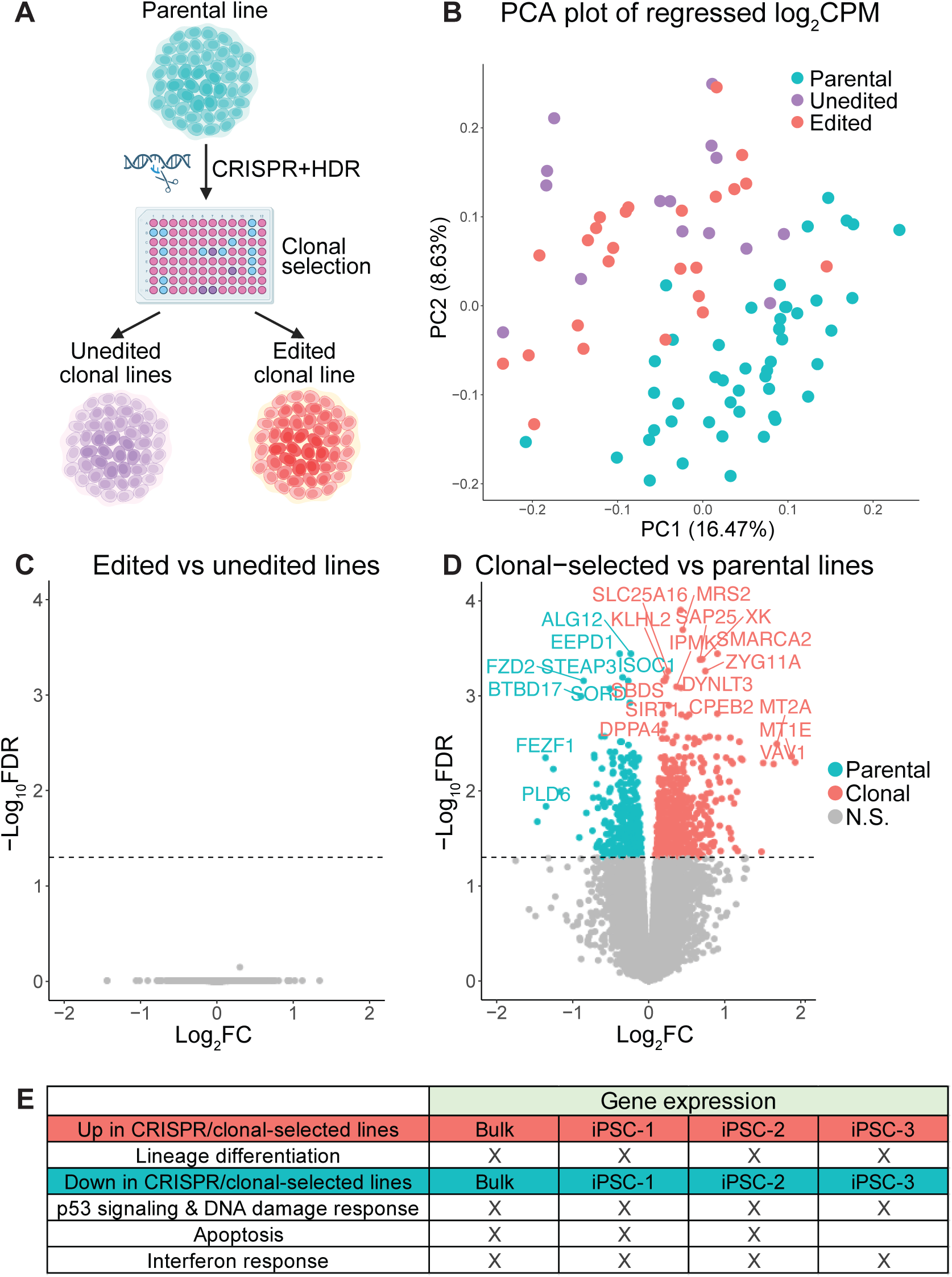
CRISPR editing & clonal selection alters p53 signaling and cholesterol metabolism (A) Schematics of the CRISPR-mediated genome editing and clonal selection procedure to generate isogenic lines with intended genetic changes (edited), lines undergone the same editing process but without intended genetic changes (unedited). (B) Distribution of regressed gene expression values (log_2_CPM) on the first 2 principal components. (C and D) Volcano plots of gene expression changes between edited and unedited lines (C) and between CRISPR-edited/clonal-selected lines and parental lines (D). (F) Pathways consistently changed in GSEA results across RNA-seq results (bulk and iPSC clusters) comparing CRISPR-edited/clonal-selected lines and parental lines. See also Figure S4 and Table S4. Student’s t test was used for panel E and H. See also Figure S5.

## DISCUSSION

Human iPSCs provide unprecedented opportunities for studying disease-modifying variants in relevant tissue types through directed differentiation^1–13,96^. However, the variability of genetic background, cell state and the efficiency of differentiation towards cell type-of-interest poses significant challenges for experimental design and result interpretation. Through our comprehensive molecular characterization of this cohort of commonly used iPSC lines with various *MAPT* mutations, we examined both inter- and intra-clonal heterogeneity to inform future line selection and experimental design. Using scRNA-seq, we identified 3 iPSC clusters consistently present in all of the lines, in addition to a poised iPSC cluster, whose proportion varied across lines and reflects an orthogonal aspect of iPSC quality, which is not captured by conventional QC measurements^97–99^ (Figures 2A and 2B). We observed a formative to primed pluripotency gradient, progressing from iPSC-1 to iPSC-3 to iPSC-2, with the poised cluster representing the least pluripotency state. We nominated a list of surface protein markers, the combination of which could be used to examine the proportion of cells in the poised iPSC cluster, as well as potentially to purify and functionally characterize each subcluster (Table S2C). In future work, it would be of value to identify whether a hierarchical relationship exists across the subclusters and whether they differ in proliferation, pluripotency and differentiation capacity.

We detected few *de novo* pPAVs in ND/NDD or constrained genes, with the exception of recurrent 20q11.21 duplications, which were associated with passage number, as previously observed^41,42^. We nominated *ID1* as a modifier that is likely driving increased resistance to forming well-patterned neural organoid, which is supported by recent over-expression studies^84^. Ideally, iPSC lines from earlier passage should be preserved and subsequent passages should be screened routinely for common chromosomal abnormalities. However, this can be logistically challenging. Alternatively, we showed that higher SB431542 concentration is a worthwhile mitigation strategy to improve neural organoid formation efficiency in lines with 20q DUP (Figure 4F). Overall, these data support the quality and genomic integrity of the majority of the lines in our study.

We observed that lines that form poorly patterned neural organoids may have different underlying causes, therefore necessitating tailored mitigation strategies. As discussed previously, lines with 20q DUP might require titration of SB431542 concentrations to overcome its effects on neural differentiation (Figures 4C and 4F). In contrast, a subset of the lines, in particular FA14530.1 lines, had disproportional percentage of the poised iPSC clusters. As we have shown previously, these lines can benefit from using slow-releasing FGF2-DISC to improve pluripotency and organoid patterning^57^. Furthermore, we showed that regardless of 20q duplication status, targeting protein translation and mitochondrial function pathways were predicted to reverse gene expression changes and potentially improve organoid formation in poorly patterning lines, which although supported by other published observations^86,100–102^, should be confirmed in future studies.

By comparing iPSC clones that have undergone CRISPR-mediated genome editing to parental lines, we observed a significant reduction in p53/DNA damage signaling and the interferon response pathway, even though we did not detect apparent mutations in these genes. The reduction of these signaling pathways most likely reflects a survivorship bias, where clones that down-regulated these proliferation-inhibitory pathways tend to out-compete others and form viable colonies. These results highlight the importance of using isogenic controls that have undergone the same genome editing procedures as clones with introduced mutations, which showed minimal transcriptional changes in this study (Figures 6B and 6C). Relying solely on parental lines as control to study the effects of targeted mutations might be cofounded by genome-editing status, even when passage numbers are controlled for.

Overall, our in-depth genomic analysis underscores the robustness of this widely used cohort of iPSCs for neuroscience research. This study establishes a rigorous framework and demonstrates the value for the genomic benchmarking of new and existing iPSC lines, thereby streamlining quality control and bolstering the reliability of downstream data interpretation. Based on these results, we recommend supplementing standard quality control steps with the following measures: 1) WGS with 30X coverage is recommended for a new founder line, as we observed that most of the putative protein-altering variants were likely germline (Figures 3D, 3E, S3I and S3J). Subsequently low pass WGS or qRT-PCR can be conducted at regular intervals (every 5-10 passages) to control for structural variants that may arise as cells are passaged, such as 20q DUP^75^. 2) If 20q DUP is already present in the line and neural differentiation is impaired, titrating SB431542 concentrations could improve differentiation efficiency. 3) When screening for clones after CRISPR editing, we recommend to sequence sufficiently long segments of the target region to include heterozygous SNPs, as suggested by Slimkin *et al*.^111^. This guarantees biallelic amplification of the locus, since large complex editing by Cas9 may remove primer binding sites (Figure 3C). 4) When choosing isogenic controls for genome-edited lines, it is strongly recommended to use lines processed in parallel during CRISPR-editing, rather than parental lines. If such controls are not available, caution should be applied when interpreting gene expression changes related to cell survival, p53/DNA repair pathways and interferon signaling pathways (e.g. Figure 6E). We submit that these steps should improve interpretability and robustness of data in this rapidly expanding field.

## Limitations of the study

Our analysis of 85 unique human iPSC lines represents the largest collection of systematic multiomic profiling of human iPSCs of which we are aware to date. Nevertheless, there remain limitations. The characterization of neural organoid formation efficiency and the effects of higher SB431542 concentration were limited to 22 and 28 lines, respectively. The smaller sample size reduced our statistical power and made it harder to control for potential confounding covariates. Additionally, we restricted our WGS analysis to germline rather than somatic variants due to the inaccessibility of donor DNA before reprogramming. As a result, we might have missed *de novo* variants present in subclones with low frequency. However, if a *de novo* variant confers a strong selective advantage, such as with 20q DUP, we should be able to identify them. Therefore, it is unlikely that we dramatically underestimated the frequency of undesirable *de novo* mutations in most of the lines.

## Star Methods

### Resource Availability

#### Lead Contacts

Further information and requests for data, resources and reagents should be directed to and will be fulfilled by the Lead Contact, Daniel Geschwind and Sally Temple.

#### Materials Availability

iPSC lines used in this study are available for request from the Tau Consortium cell line collection (https://www.neuralsci.org/tau)^18^.

#### Code Availability

All the code used in this study will be available on Github upon publication, and the pipelines described in the Method Details.

### Experimental Model and Subject Details

#### Cell lines

This study included 85 iPSC lines from 25 different donors (Figure S1 and Table S1) obtained from the Tau Consortium cell line collection (https://www.neuralsci.org/tau);^1^ All lines were negative for mycoplasma, and karyotypically normal by G-banded Karyotype (WiCell Research Institute, Inc.). The iPSCs were maintained in six-well plates coated with growth factor-reduced Matrigel or Cultrex at 37°C and 5% CO_2_. The cultures were fed with twice a week mTeSR1 medium with an FGF2-DISC unless otherwise specified. One FGF2-DISC was added into one well of a 6-well plate with 2 mL of mTeSR1 medium for approximately a week. The medium, but not the FGF2-DISC, was replaced every 2-3 days based on culture confluency. In either culture method, iPSC cultures were clump-passaged using ReLeSR about once a week. Cells were not allowed to grow past 80% confluency. To collect cells for genomic profiling, iPSC colonies were digested to single cells using Accutase or TrypLE Express. Samples without FGF2-DISC were only included for initial joint processing, such as joint calling in WGS and clustering for scRNA-seq analysis, except in Figures 2B-2D and S2B-S2M.

## Method Details

### Organoid production

Organoids were generated at the NeuraCell core facility (Neural Stem Cell Institute, NY, USA). When iPSC cultures reached about 70-80% confluency, the medium was aspirated, and the wells were rinsed twice with DMEM/F12. Following, 1.5 mL of TrypLE Express or Accutase was added per well of the 6-well plate and incubated for 7-10 minutes at 37°C, 5% CO_2_ until cells detached from the dish. Using a 1000 µL pipette, gentle trituration was performed to achieve a single cell suspension. The cell suspension was transferred to a 50 mL conical tube, and cells were washed with DMEM/F12 two times. Following, cells were counted manually with a hemocytometer and resuspended at 1 million cells/mL in mTeSR1 with 10 µM ROCK inhibitor Y-27632 (Tocris). To set up organoids in one 96-slitwell plate (S-bio, MS9096SZ), 0.5-1 million cells in 1 mL were added to 9 mL mTeSR1 plus 10 µM rock inhibitor. 100 µL of this suspension was dispensed into each well to yield 5-10,000 cells per well, with the goal of achieving spheroids of 350-500 µm diameter. The 96-well plate was incubated at 37°C and 5% CO_2_ overnight to generate spheroids. The next day (day 0 of differentiation), 20 mL DMEM/F12 medium was added, the plate was gently rocked to wash. This wash with gentle rocking was repeated a second time. The medium was then replaced with 14 mL differentiation Medium A: E6 medium supplemented with 2.5 µM dorsomorphin (DM), either 10 or 20 µM SB431542 and 2.5 µM XAV939. Lyophilized vials of each small molecule were freshly reconstituted and aliquoted for each organoid batch.

Plates were fed daily by slightly tilting the plate, gently aspirating about 14 mL of the pooled medium and replacing with freshly made Medium A, achieving approximately 65% medium exchange from day two until day five. On the sixth day, the medium was changed to neural medium (NM), consisting of Neurobasal-A plus B-27 supplement without vitamin A, GlutaMax and Anti-A and supplemented with 20 ng/mL EGF plus 20 ng/mL FGF2. NM plus EGF/FGF2 (Medium B) was changed daily for 10 days then every other day (3X/week) for 9 days with 65% media exchanges. On day 25, the medium was replaced with NM supplemented with 20 ng/mL BDNF and 20 ng/mL NT3 (Medium C) with 65% medium feeds every other day (3X/week). From day 43 onward, the medium was changed every other day (3x/week) using NM without added growth factors with 15-20 mL per dish (75% medium changes.)

Organoids were harvested for QC on day 20 and at 2 months and assessed by qPCR and immunohistochemistry (IHC) of organoid sections. Cortical markers were evaluated by IHC: PAX6 and FOXG1 at day 20 and CTIP2/BCL11B and TBR1 at 2 months. Efficiency of organoid formation was defined as positive FOXG1 staining at day 20, which strongly correlates with correct organoid patterning in later time points.

### Fixation and frozen sectioning of organoids

Organoids were fixed using 4% paraformaldehyde at 4°C for 2 hours for 20-day timepoints or overnight for older organoids. They were then rinsed three times in PBS and allowed to sink in 30% sucrose in PBS overnight. The organoids were placed in cryotrays (Seal N Freeze) with OCT compound (Tissue Tek, 4583), snap frozen in a slurry of dry ice and isopropyl alcohol in a Seal N Freeze box and stored at -80°C. Organoids were cryostat-sectioned sequentially at 20 µm thickness using a Leica cryostat (model CM3050S). Sections were placed on microscope glass slides, dried overnight, and stored at -20°C for subsequent immunohistochemistry.

### Immunohistochemistry staining for organoid sections

Slides were thawed, brought to room temperature (RT), and lines drawn to partition the edges of sections on the glass slides using a PAPpen. The sections were rehydrated, blocked, permeabilized and immunostained with primary antibodies: PAX6 (1:10-1:100), FOXG1 (1:500), CTIP2 /BCL11B (1:500), SATB2 (1:50), BTUB III (1:1000), MAP2AB (1:1000-1:2000), TBR1 (1:500). Primary antibodies were incubated overnight at 4°C, washed three times with PBS and then incubated with corresponding Alexa Fluor conjugated secondary antibodies (1:333-1:1000) for 1 hour at RT. Sections were coverslipped and imaged using fluorescence and confocal microscopy (Zeiss AXIO Observer.Z1; Zeiss 780). See Key Resource Table for antibody information.

### Flow cytometry of iPSCs

When iPSC cultures reached about 80% confluency, the medium was aspirated and wells rinsed twice with DMEM/F12. 1.5 mL of TrypLE Express or Accutase was added per 6-well and incubated for 7-10 minutes at 37°C, 5% CO_2_ until cells detached from the dish. Using a 1000 µL pipette, gentle trituration was performed to achieve a single-cell suspension, which was then transferred to a 15 mL conical tube. Cells were washed with DMEM/F12 two times. Following, cells were resuspended in 1mL of PBS without calcium and magnesium (PBS-/-) and 9 mLs of ice-cold methanol was slowly added while vortexing. On the day of staining, fixed cells were washed three times with PBS-/-. Cells were then counted using a hemocytometer and resuspended in 3% BSA/PBS-/- at 1 million cells per 100 µL. One million cells were incubated with TRA-1-60 (5 µL/ million cells), SSEA4 (20 µL / million cells), and SSEA1 (20 µL / 1 million cells) antibodies for 30 minutes at RT. After 2 washes with PBS-/-, stained cells were analyzed using a BD FacsAria II Flow Cytometer/Cell Sorter. Single-stained and unstained controls were used to set the gates. See Key Resource Table for antibody information.

### iPSC RNA extraction and bulk RNA-seq library preparation

RNA was extracted from samples using the Zymo Research Quick-RNA Miniprep Plus kit. The concentration was measured with Qubit RNA High Sensitivity Assay kit. The RNA Integrity Number (RIN) was evaluated for each sample on an Agilent TapeStation (all RIN > 9). Library preparation was completed using the Illumina TruSeq Stranded Total RNA kit. 100 or 150 bp paired-end sequencing was completed on an Illumina NovaSeq 6000 or NovaSeq X plus to acquire around 50 million fragments per sample. See Key Resource Table for reagent information.

### iPSC RNA-seq preprocessing and differential expression analysis

Adaptor-trimmed RNA sequencing files were mapped to the genome (GRCh38; https://ftp.ensembl.org/pub/release-106/fasta/homo_sapiens/dna/ Homo_sapiens.GRCh38.dna.primary_assembly.fa) using RNA-STAR^103^ (v2.7.10a). Gencode annotations (Release 40, GRCh38.p13) were used from ENSEMBL. Samples sequenced on different lanes were merged using Samtools^104^ (v1.15). Sequencing metrics were obtained using Picard tools (2.27.3)^105^. Finally, RSEM^106^ (v1.3.3) was used for quantification to obtain expected read counts at the gene and transcript levels.

Raw gene read counts were then subjected to several processing steps in preparation for downstream analysis, mainly using R (v4.3). Genes were first filtered with filterByExpr() function in edgeR^107^ (v4.0.16) using default settings, unless otherwise specified. Genes with an effective length (measured by RSEM) of less than 15 bp were also removed. The counts for the remaining genes passing these filters were normalized using edgeR with an adjustment for sample read depth variance. An offset value calculated with CQN (v1.48.0)^108^ accounting for GC content bias and gene effective length bias in read quantification was also incorporated during the normalization process.

To identify differentially expressed genes, a linear mixed-effect model (LMM) was used to account for variations across different iPSC donors or batches as random effects variables with variancePartition^109^ (v1.32.5) R package. Top 5 principal components were calculated using sequencing metrics (SeqPCs) generated by Picard tools and included in the final models along with Sex and RIN unless otherwise specified. voomWithDreamWeights(), dream() with Kenward-Roger approximation and eBayes() functions were used to extract log_2_FC and FDR for each gene. Regressed log_2_CPM values were calculated using residuals and coefficients calculated by fitVarPartModel(). Variance partitioning was done using fitExtractVarPartModel().Reanalysis of SMAD2 and SMAD3 over-expression in Nakatake *et al*.^83^ was done using the same method as described above.

### Gene set enrichment analysis (GSEA)

fGSEA (v1.28.0)^111^ was used with a pre-ranked gene list by t-statistics obtained from previous steps. Pathways obtained from curated gene sets (c2.cp.v2024.1.Hs.entrez.gmt) with minimum size of 30 and maximum size 500 were included in the analysis. Only pathways with FDR<0.05 were retained and included in Table S4.

### Rank Rank Hypergeometric Overlap (RRHO) analysis

RRHO (v1.42.0)^112^ R package was used to generate p-values and correlation heatmaps. To compare two DEG lists, common genes were selected and log_2_FC values were used to perform RRHO with step=200 and alternative = “two.sided” to account for negative correlations.

### iPSC DNA extraction and short-read WGS

DNA was extracted from samples using the Zymo Research Quick-DNA Miniprep Plus kit. The concentration was measured with Qubit DNA High Sensitivity Assay kit. Initial WGS libraries were generated using PCR-free NEBNext^®^ Ultra™ II DNA Kits and 150 bp paired-end sequencing was done on an Illumina NovaSeq 6000 at Novogene. Supplemental libraries were generated using PCR-free NEBNext^®^ Ultra™ II FS DNA Kits in-house and 150 bp paired-end sequencing was done on an Illumina NovaSeq X plus to get 30x coverage total.

### WGS preprocessing and variant calling

Adaptors were first trimmed from raw FASTQ files using cutadapt (v4.1)^113^. Trimmed reads were used as input for DRAGEN-GATK (v4.5.0.0)^114^ Maximum Quality mode for preprocessing and the germline short variant discovery pipeline^115^ to generate individual gVCF files. Subsequently, joint genotyping pipeline was used to generate cohort short variant VCF file. Downstream processing was done using BCFtools (v1.11)^116^. Only “PASS” variants were retained and multiallelic sites were split. Individual genotypes with GQ<20 and DP<10 were marked as missing, and monomorphic sites or F_MISSING>0.1 were removed. VEP (v111)^117^ was used to predict the effects of variants on protein functions. In parallel, BAM files generated from previous alignment steps were fed into GATK-SV (v1.0.1)^118^ joint calling pipeline for SV calling and QC plotting. Only “PASS” and “MULTIALLELIC” variants were retained for downstream analysis. Individual MULTIALLELIC genotypes with CNQ<20 were marked as missing, and monomorphic sites or F_MISSING>0.1 were removed. Annotations from GATK-SV pipeline was used to predict SV effects on protein functions.

### WGS downstream analysis

Rare variants (< 0.01%) in short variant VCF file were identified by removing iPSC variants overlapping with variants in gnomAD v4.1 with AF_grpmax_joint > 0.0001. Similarly, rare SVs were identified by removing iPSC SVs overlapping with SVs in gnomAD v4.1 with AF>0.0001 in any populations or equivalent variants in nstd186 (NCBI Curated Common Structural Variants). SVs were considered equivalent if both ends are within 1kb from each other using SURVIVOR (v1.0.7)^119^. Rare SVs likely to be artifacts were filtered by overlapping with non-“PASS” sites in gnomAD, as well as rare variants that are present in all the lines derived from more than one donor. pPAVs were defined as short variants annotated as HIGH impact and high confidence (HC) tag for protein-coding genes by VEP. Out-of-frame INDELs in the same gene that neutralize each other were removed. For SVs, sites predicted by GATK-SV pipeline as “MSV_EXON_OVERLAP”, “BREAKEND_EXONIC”, “COPY_GAIN”, “INTRAGENIC_EXON_DUP”, “LOF” or “PARTIAL_EXON_DUP” were treated as pPAVs. The list of constrained genes was obtained from gnomAD v4.1 with LOEUF score < 0.6 as suggested by the gnomAD team. The list of neurodegeneration-related genes was a union of genes associated with Alzheimer’s disease, amyotrophic lateral sclerosis, Lewy-body dementia or Parkinson’s disease in The Neurodegenerative Disease Knowledge Portal (NDKP). The list of neurodevelopmental disorder-related genes was derived from a combination of genes labeled as definitive by SysNDD database^120^ and genes with organ specificity labeled as brain or multisystem by DDG2P^121^ (v5.0). To estimate the number of variants per individual in non-Finish European (NFE) population in gnomAD v4.1, we calculated the allele frequency of each variant within NFE individuals and calculated the sum of all variants within each category.

For reanalysis of KOLF2.1J WGS results^62^, raw FASTQ files were obtained from AD workbench and processed the same way as the iPSCs in this cohort, except that no joint genotyping was performed for short variants and single sample mode was used for the GATK-SV step.

### Targeted long-read DNA sequencing of the *MAPT* locus

Library preparation and enrichment of MAPT locus was done at The Center for Advanced Genomics Technology at Icahn school of medicine at Mount Sinai using custom-designed capture probes and library prep kit from Twist Bioscience. Libraries were sequenced on PacBio Sequel II (batch 1) or Revio (batch 2) and raw data were preprocessed to generate HiFi BAM files at Mount Sinai. Reads were aligned to GRCh38_no_alt_analysis_set.fasta using pbmm2 (v1.17.0). Deepvariant (v1.6.1)^123^ and GLnexus (v1.3.1)^124^ was used to joint call short variants. SVs were called using both pbsv (v2.9.0) and Sniffles (v2.6.0)^125^, and the union was used for downstream analysis. SVs were filtered for “PASS” sites, F_MISSING < 0.1 and annotated with VEP.

### DNA methylation array, preprocessing and differential methylation analysis

DNA methylation analysis was done at UCLA Neuroscience Genomics Core (UNGC) using Illumina 850K Infinium MethylationEPIC v1.0 kit and DNA extracted from iPSCs. Raw data were imported into R using wateRmelon (v2.8.0)^126^ and outlier probes were filtered with pfilter() function. The intersection of the remaining probes from each array were merged for downstream processing. Cross-reactive and polymorphic probes identified in Pidsley *et al*^127^. and McCartney *et al.*^128^ were removed. SNPs and sex chromosomal probes present in the EPIC array were subsequently removed. Beta values were filtered and normalized using champ.filter() and PBC method of champ.norm() functions from ChAMP (v2.32.0)^129^ package respectively. Normalized beta values were then converted to M values for downstream variance partitioning and differential methylation analysis similar to bulk RNA-seq analysis. The only differences were that RIN was not included in the model and voomWithDreamWeights() step was skipped.

### Gene Ontology (GO) analysis for differentially methylated genes

gProfileR2 (v0.2.3)^130^ package was used for gene ontology enrichment. ‘BP’ (biological process) terms within DAVID database^131^ BP_FAT, KEGG pathways, Reactome pathways and Wikipathways were included in the analysis. Significant sites (FDR<0.05) were first grouped into promoters, enhancers and gene bodies using annotations from Bizet *et al*.^132^ and the corresponding genes were used as input for GO analysis. Results were first filtered for minimum and maximum term size of 30 and 500 respectively and p-values were adjusted with Benjamini-Hochberg procedure.

### Generation and sequencing of scRNA-seq libraries

Cryopreserved iPSCs were thawed and washed with PBS twice. Cells were fixed on ice with Scale FixKit (Scale Biosciences), counted and loaded into 3 wells of ScaleBio RNA kits per sample and libraries were prepared according to manufacturer’s protocol. Final libraries were sequenced on either Illumina NovaSeq 6000 or NovaSeq X plus. Raw BCL files were processed using ScaleBio RNA workflow (v1.3) to generate BAM files, count matrix and QC reports.

### Processing of scRNA-seq data

Raw count matrix was first processed with CellBender (v0.3.2)^133^ to remove ambient RNA reads. The count matrix were first filtered by scDblFinder (v1.18.0)^134^ to remove doublets and further filtered for high-quality cells (nGene > 500, nGene < 15000, %MT < 10, nUMI > 2000 and nUMI < 100000) using Seurat (v5.2.1)^135^. Data was further processed using NormalizeData(), ScaleData() to regress out percentage of exon reads, FindVariableFeatures(), RunPCA() and IntegrateLayers() using “FastMNNIntegration”^136^, RunUMAP(), FindNeighbors() and FindClusters() with the Leiden algorithm. The parameters for number of PCs, number of neighbors and resolution of clustering were determined using bootstrapping method with 80% of the cells bootstrapped 20 times with replacement from each sample. After repeating the processes described above for each bootstrapping iteration to get clusters, the results were compiled using scclusteval (v 0.0.0.9000)^137^. We selected 40 PCs, k=30 neighbors and resolution=0.15 as final parameters with almost 100% of the cells in stable clusters, which were defined as clusters with maximum Jaccard index > 0.6^137^. SingleR (v2.6.0)^138^ was used to map the data to Human Primary Cell Atlas^53^.

To perform DEG analysis across clusters, we used dreamlet (v1.2.1)^139^ package to generate pseudo-bulk counts of the count matrix by sample ID and clusters. Only cluster/sample ID combinations with more than 50 cells were retained. DEG analysis was done in the same fashion as for bulk RNA-seq described above. SeqPCs derived from sequencing metrics calculated by Picard tools on BAM files split by cell barcodes for each cluster of each sample. To find top cluster markers, we included sample ID and batch as random effects variables in our model and cluster as variable of interests, plus sex and top 5 SeqPCs as covariates. Genes significantly up-regulated in each cluster compared to all 3 other clusters (FDR<0.05) were retained for GO analysis described previously. To calculate gene network module (GNM) scores, we obtained the list of genes in formative module (GNM5) and primed module (GNM10) as described in Arthur *et al.*.^54^ The sums of regressed log_2_CPM for all genes in the module were used as module scores. To identify surface protein markers for each cluster, the list of human cell surface protein was first obtained from Bausch-Fluck *et al*.^140^. We used lasso regression from glmnet (v4.1.8)^141^ package with 10-fold cross-validation to select surface markers using regressed log_2_CPM values. To perform differential composition analysis, we used scCODA (v0.1.9)^142^ to identify significant changes of iPSC cluster frequencies across conditions. Reference clusters were chosen automatically by the software. Reanalysis of scRNA-seq data from Osnato *et al*.^82^ was done the same way as this iPSC dataset, except that all cells were included for pseudo-bulk aggregation. Processed results from *Feng et al.*^86^ was downloaded directly from https://figshare.com/s/14edeeab56eb8a885df3.

### Statistics

All statistical analysis and data visualization was performed in R or python. Box plots and violin plots are used to present most of the data with individual data points. Statistical tests used are specified in the corresponding figure legends. Significances are plotted as * P< 0.05; ** P<0.01; *** P<0.001.

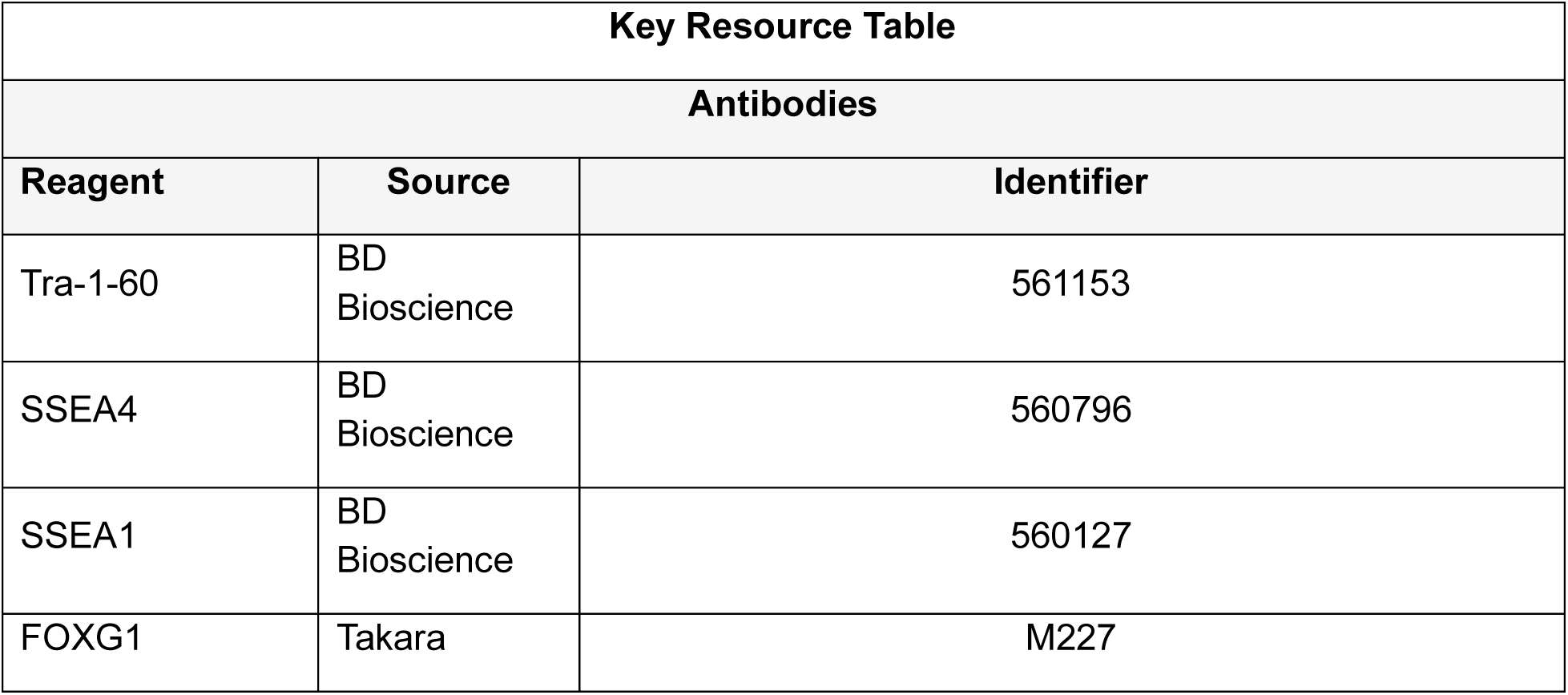
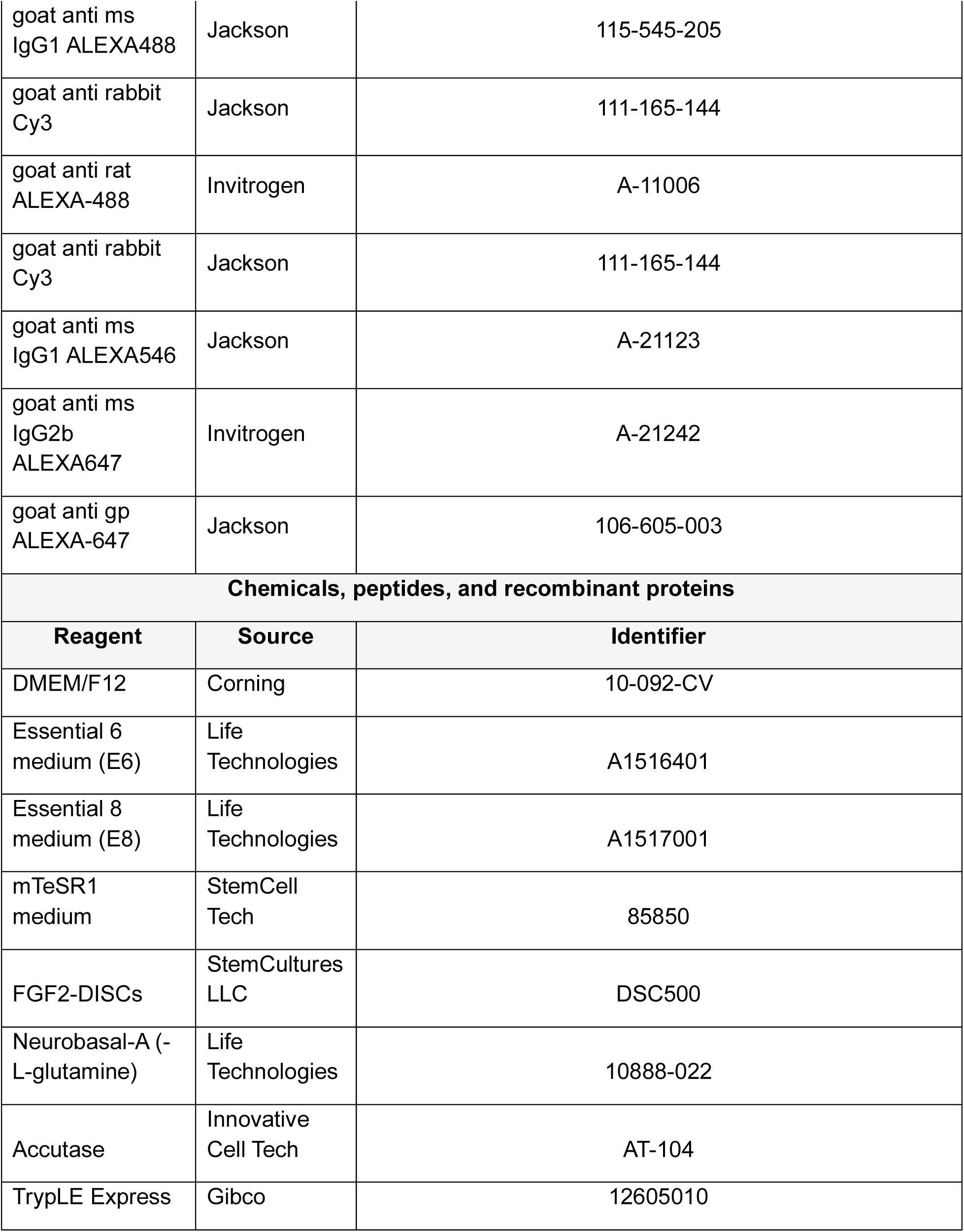
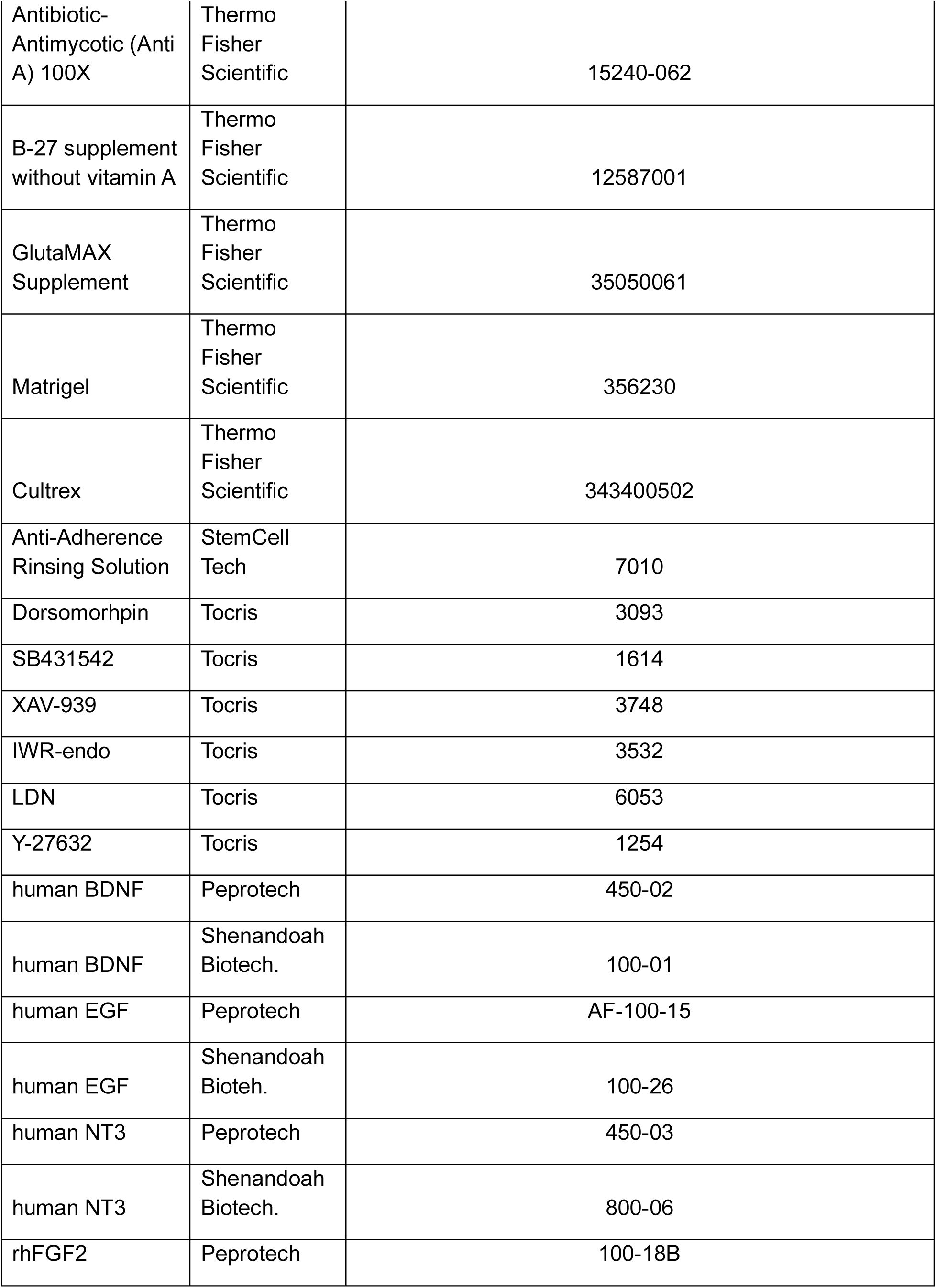
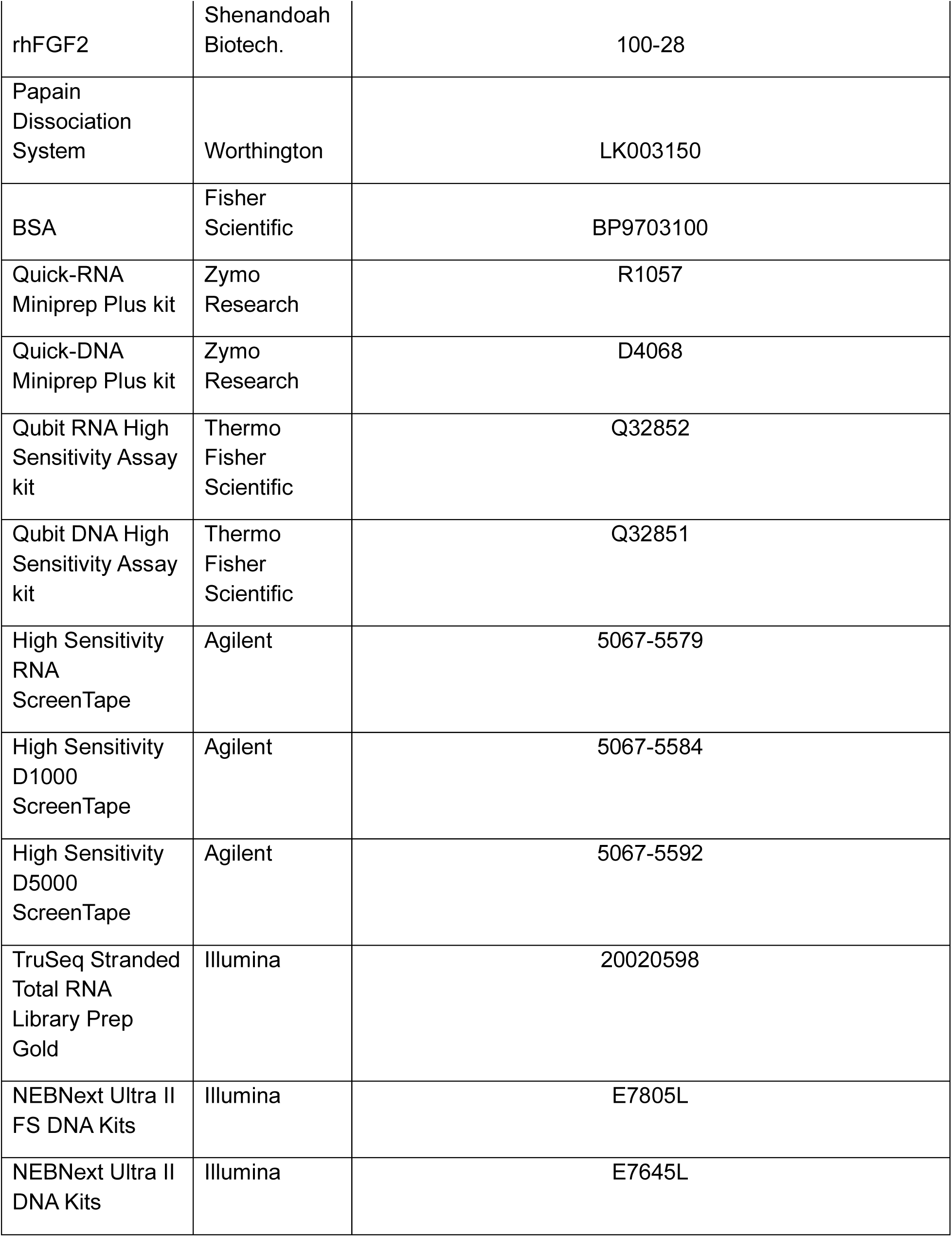
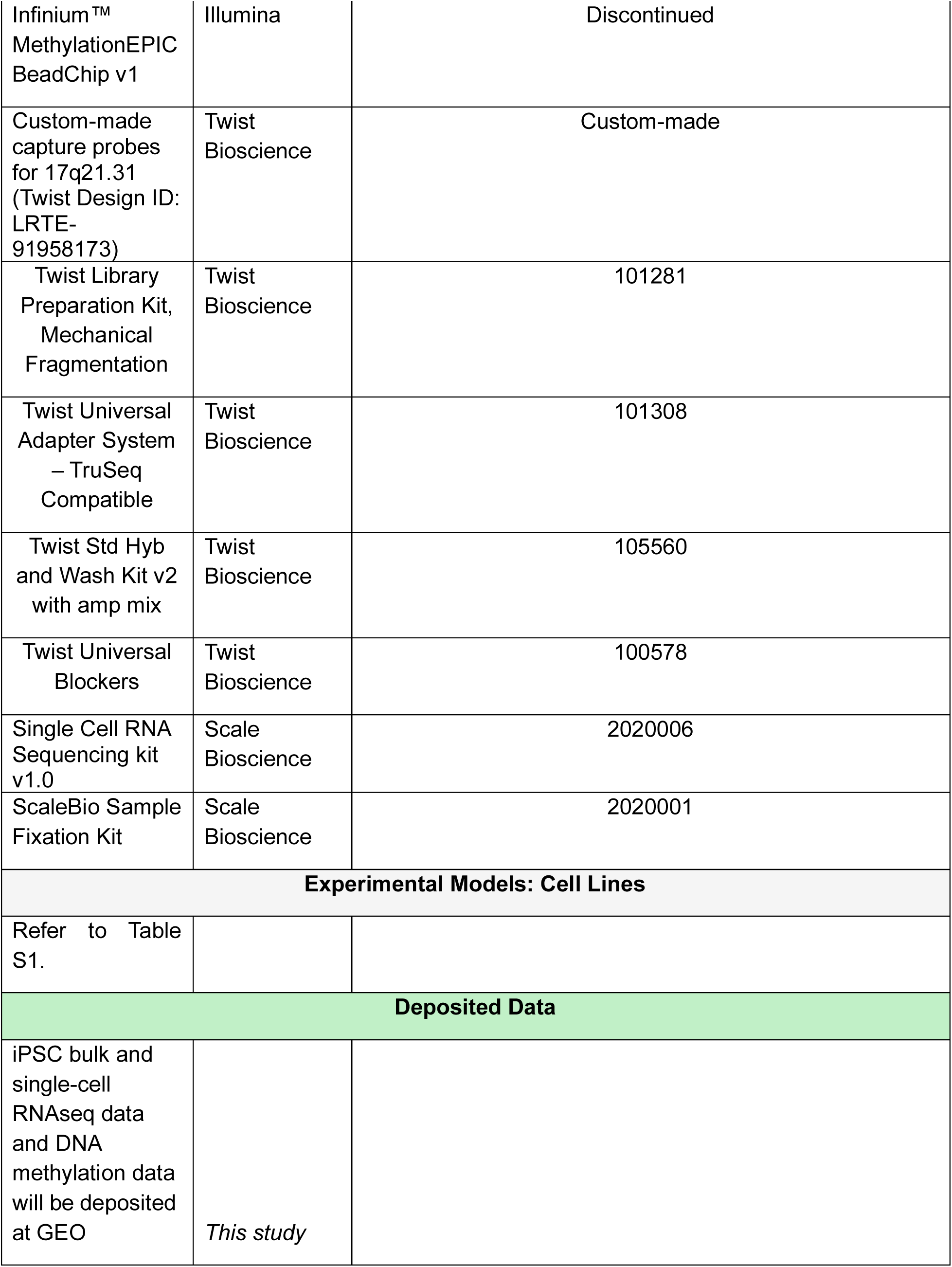
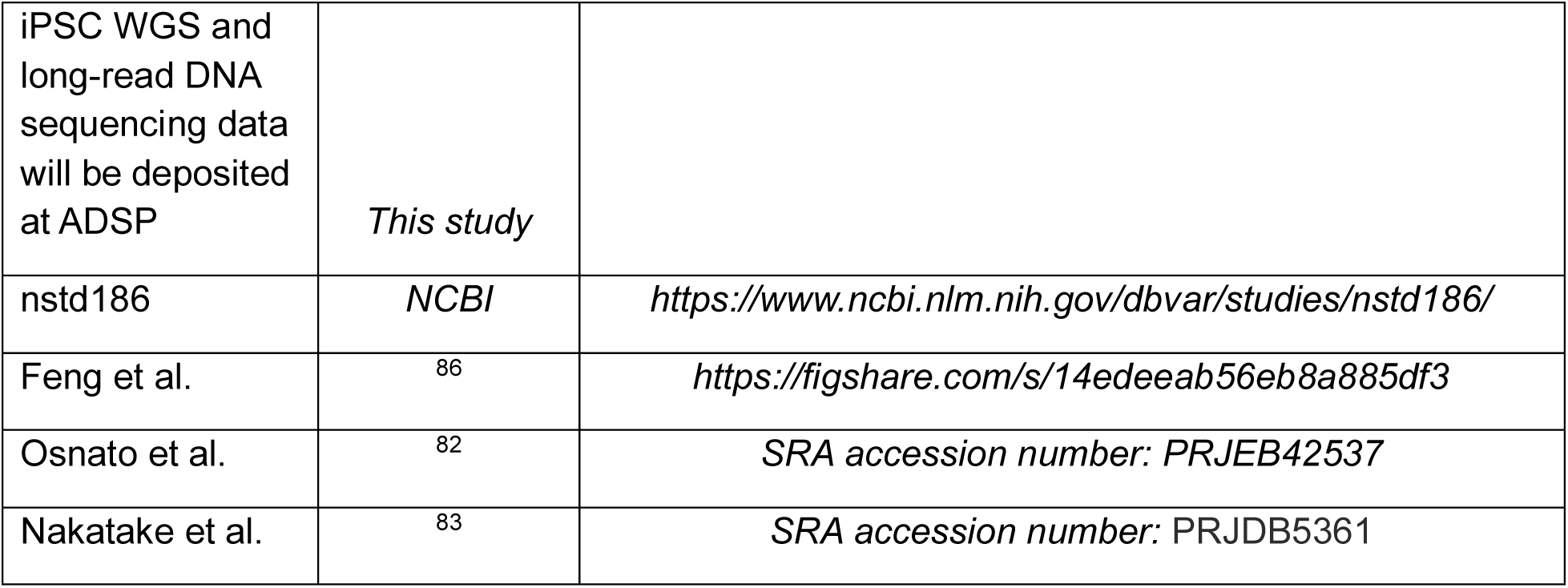
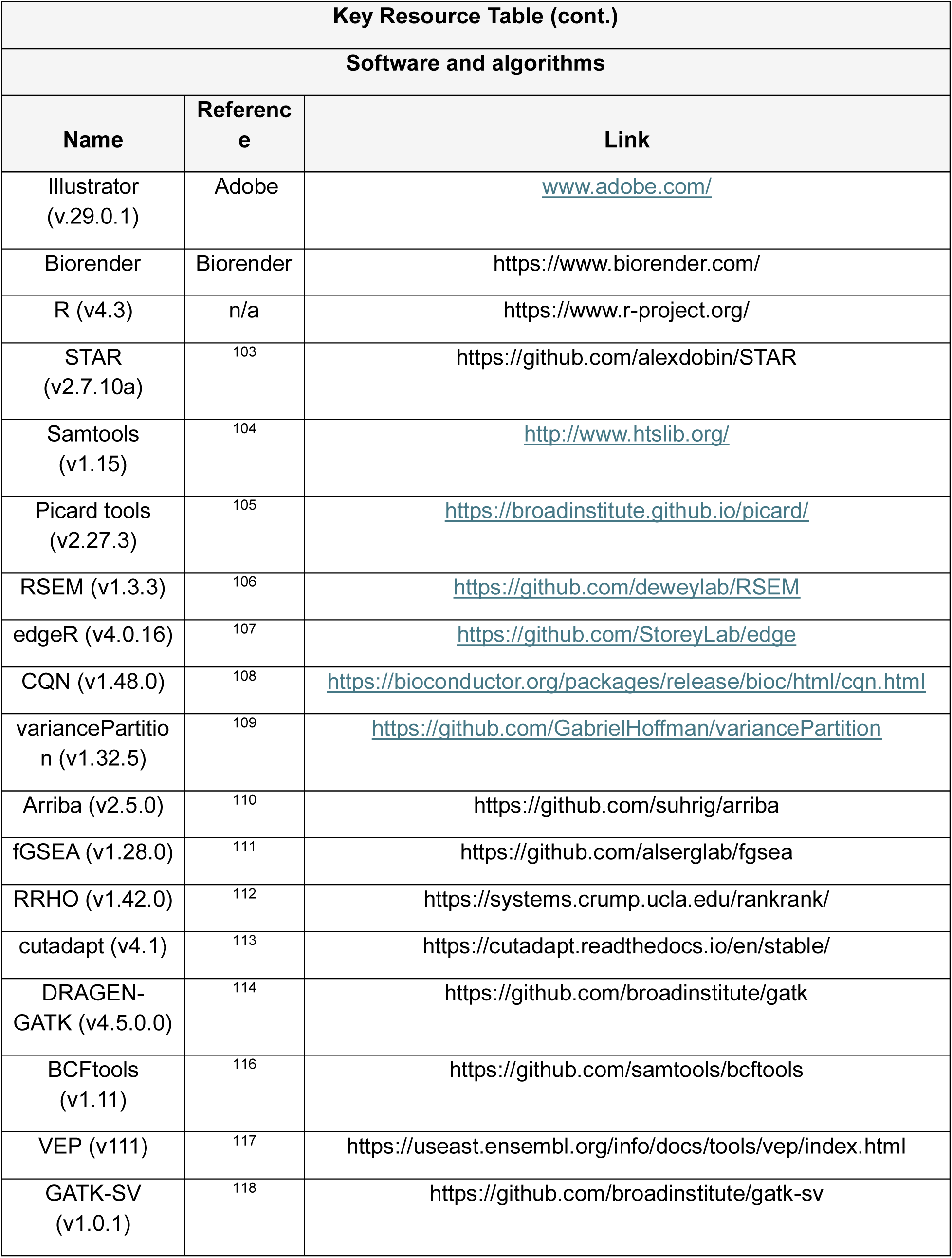
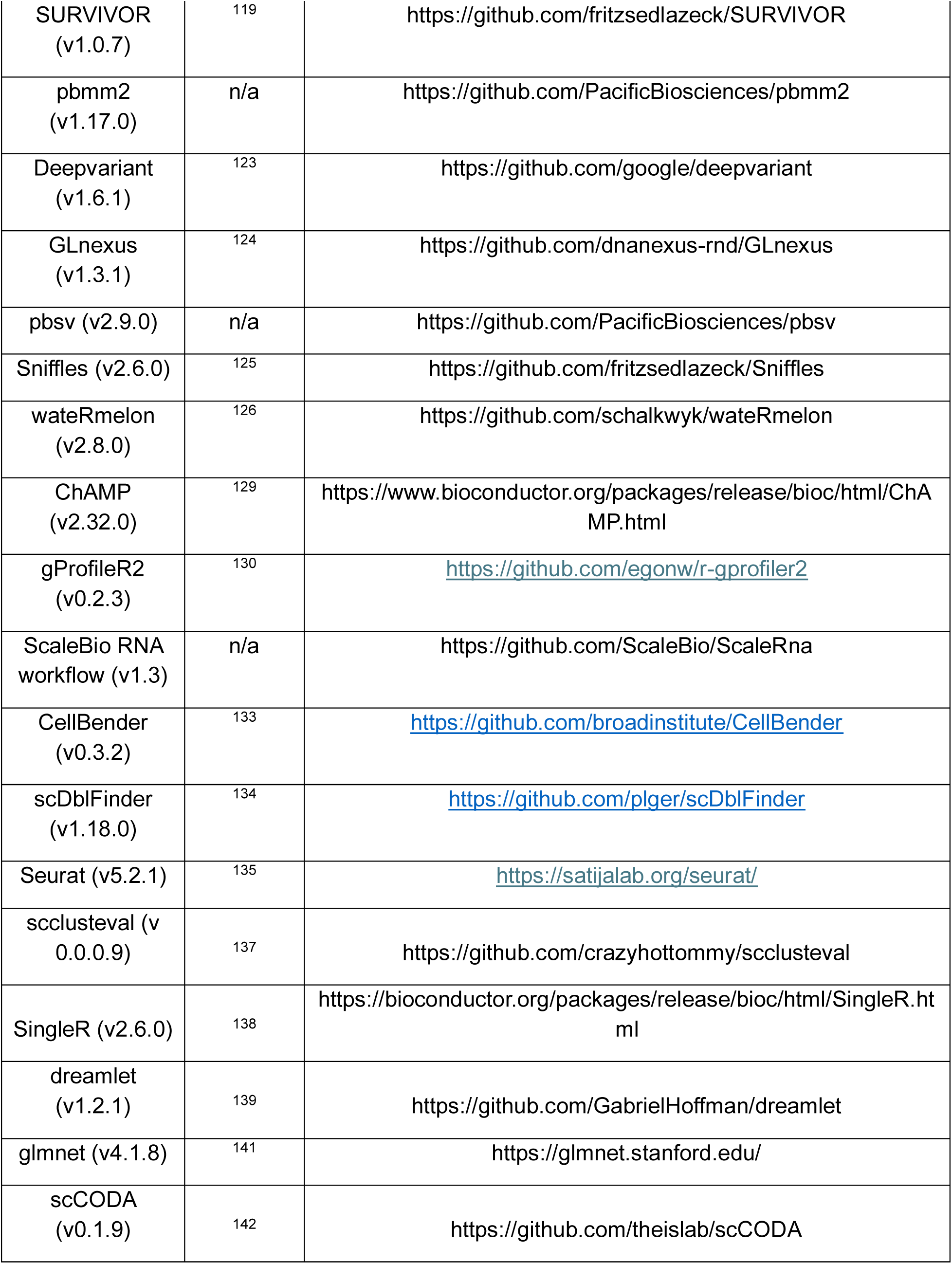
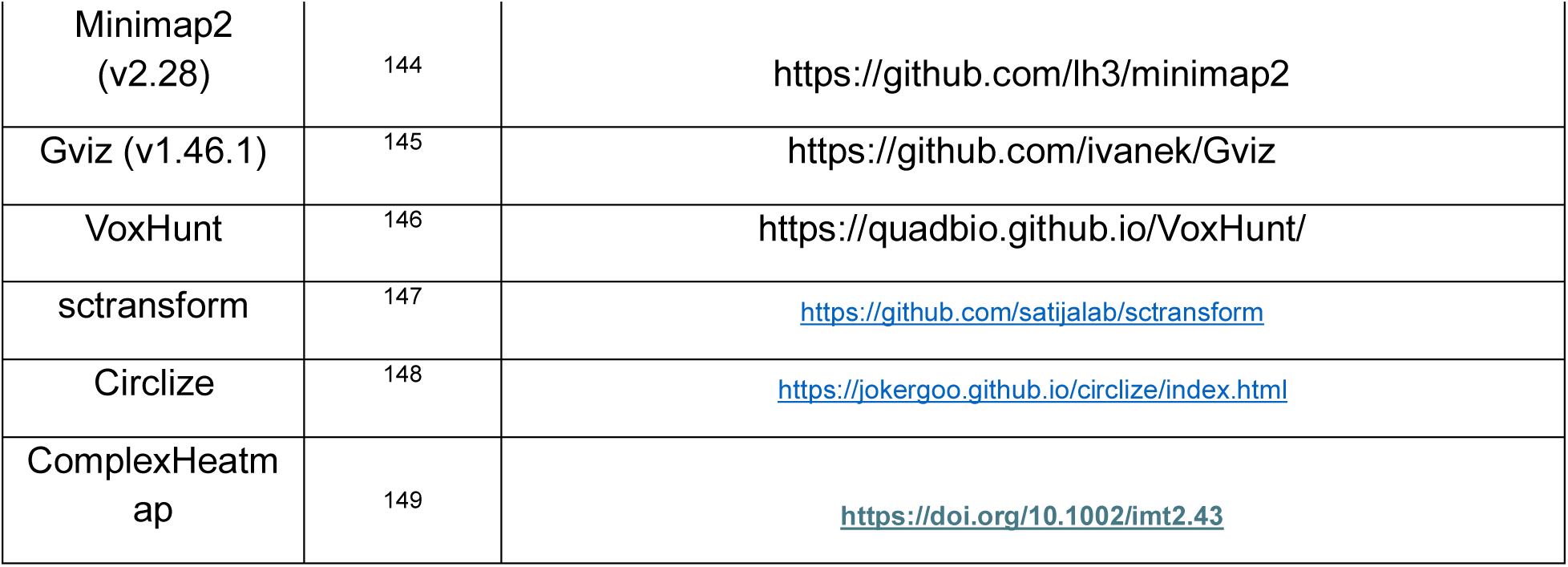

## Supporting information

Supplemental Table 2 for scRNA analysis

Supplemental Table 4 for DEG analysis

Supplemental Table 3 for WGS analysis

Supplemental Table 1 for iPSC metadata

## ACKNOWLEDGEMENTS

The authors thank members of the Tau Consortium for generating and characterizing iPSC clones used in this study. We also thank the UCLA Neuroscience Genomics Core, UCLA Broad Stem Cell Research Center Sequencing Core, UCLA Technology Center for Genomics & Bioinformatics and the Center for Advanced Genomics Technology at Icahn school of medicine at Mount Sinai for library preparation and sequencing. Computation resources were provided by UCLA Hoffman2 High-Performance Compute Cluster.

This work was supported by the National Institutes of Health grants UG3NS104095 (DHG), 5U54NS123746 (DHG), R35NS097277 (ST), RF1NS123568 (ST), R01NS142335 (ST), R01NS110890 (CMK), P30AG066444 (CMK), the Tau Consortium and Rainwater Charitable Foundation (DHG, ST, CMK).

---

**Figure S1.**
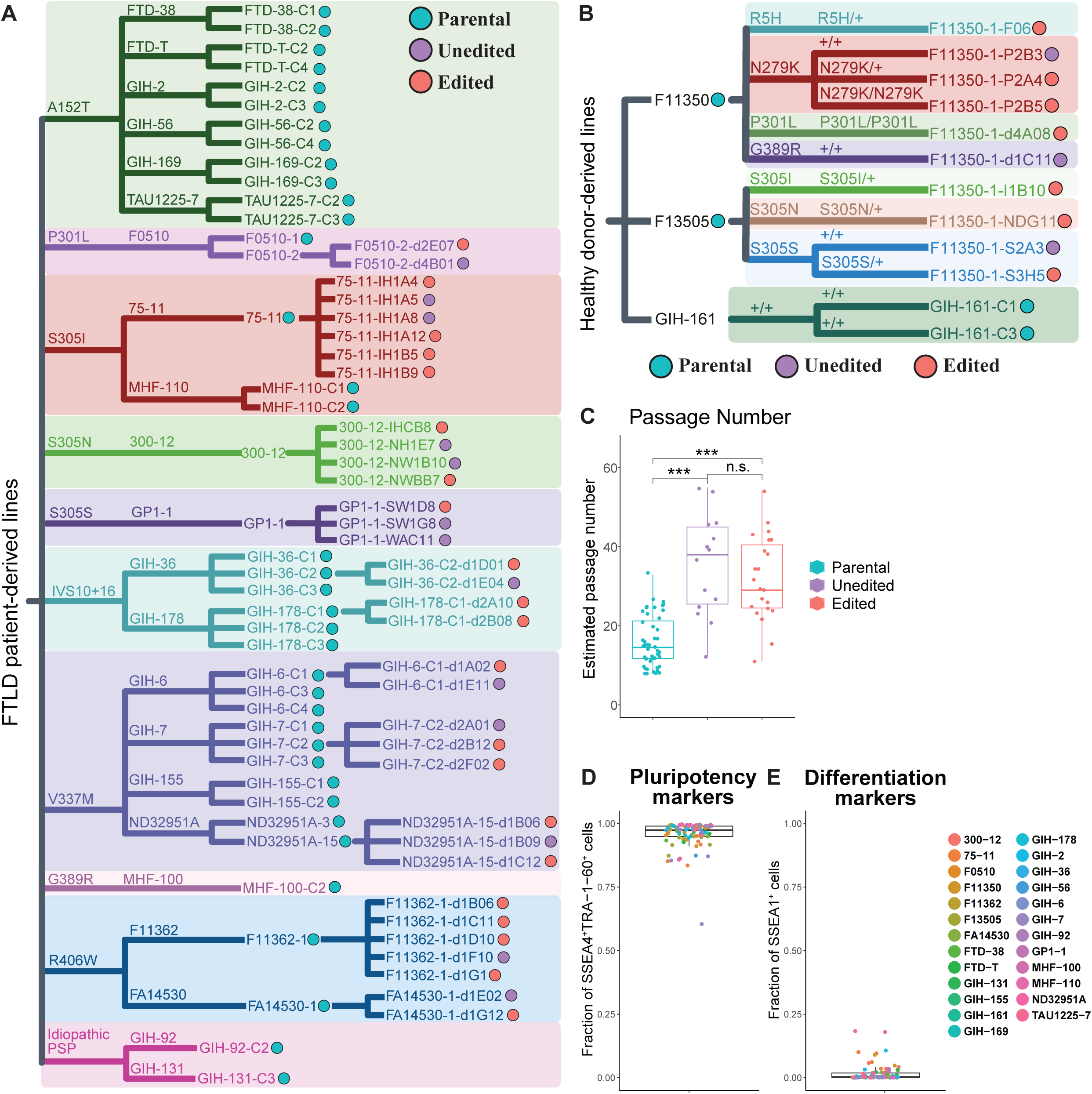
Related to Figure 1: Detailed description of the iPSC cohort in this study (A and B) Detailed list of FTLD patient-derived (A) and healthy donor-derived (B) iPSC lines grouped by MAPT mutations and donors. Each line is color-coded to represent their CRISPR-editing status. (C) Estimated passage numbers of iPSC lines grouped by their CRISPR-editing status. One-way ANOVA and Tukey’s method for post-hoc comparisons. (D and E) Fraction of SSEA4^+^TRA−1−60^+^ (pluripotency marker) cells (D) and SSEA1^+^ (differentiation marker) cells (E) as measured by flow cytometry. See also Table S1.

**Figure S2.**
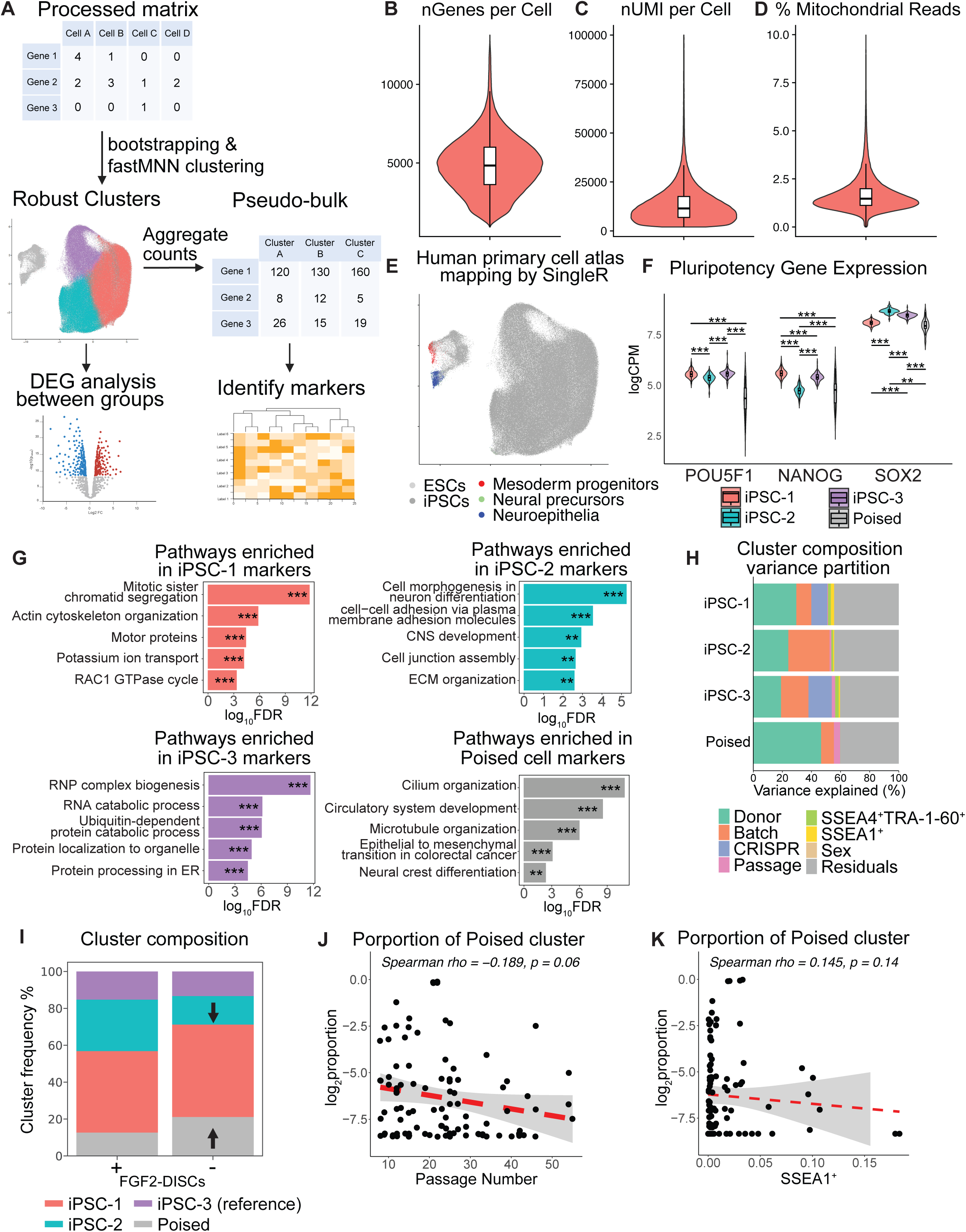
Related to Figure 2: Data associated with scRNA-seq analysis of iPSC lines (A) Schematics of our bioinformatics pipeline for scRNA-seq analysis. (B-D) Basic QC measurements of the scRNA-seq dataset, showing 4882 ± 1777 genes detected per cell (B), 13738 ± 10386 UMIs detected per cell (C) and 1.7 ± 1.0% mitochondrial reads per cell (D) after preprocessing and filtering. (E) UMAP representation of cell types in Human primary cell atlas^53^ to which the cells were mapped to using SingleR^150^. (F) Violin plot of regressed pseudo-bulk logCPM values of POU5F1, NANOG and SOX2 in different iPSC clusters. (G) Selected pathways enriched in GO analysis using genes significantly up-regulated in iPSC-1 (tope left), iPSC-2 (top right), iPSC-3 (bottom left) and poised iPSC cluster (bottom right) compared to the rest of the clusters.(H) Variance partitioning of each Leiden cluster by known technical and biological covariates using VariancePartition^56^. (I) Differential cluster composition analysis between lines grown in medium with or without FGF2-DISCs using scCODA^151^. Significant changes (FDR < 0.05) were labeled with arrows. (J and K) Correlation between the log_2_proportion of poised cluster in each iPSC line and passage number (J) or percentage of SSEA1^+^ cells (K). See also Table S2.

**Figure S3.**
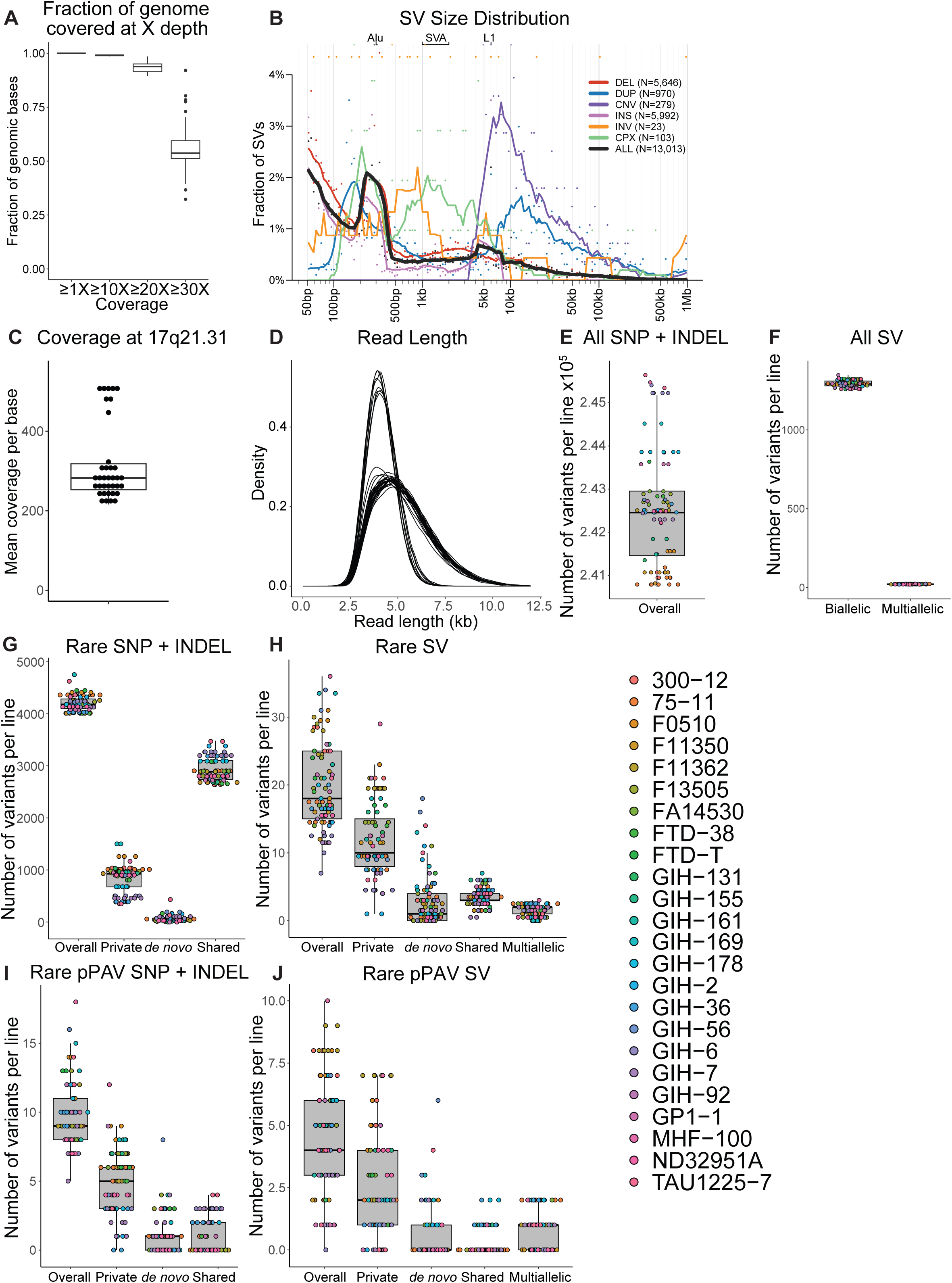
Related to Figure 3: Data associated with WGS and long-read DNA sequencing of the MAPT locus of iPSC lines (A) Fraction of genome covered at 1X to 30X by short-read WGS for each iPSC line (B) Size distribution of SVs called in WGS data grouped by SV classes (C and D) Mean coverage (C) and read lengths (D) of the 17q21.31 locus in long-read DNA sequencing data. Library generation and sequencing were done in two batches, reflecting the bimodal distributions of the coverage and read lengths. (E-J) Numbers of SNPs and INDELs (< 50bp) and SVs (>50bp) called in each iPSC line for all variants (E and F), rare variants with minor allele frequency (MAF) < 0.01% (G and H) and rare pPAVs (I and J). Variants were grouped by their frequency among iPSC lines as described in Figure 3D. See also Table S3.

**Figure S4.**
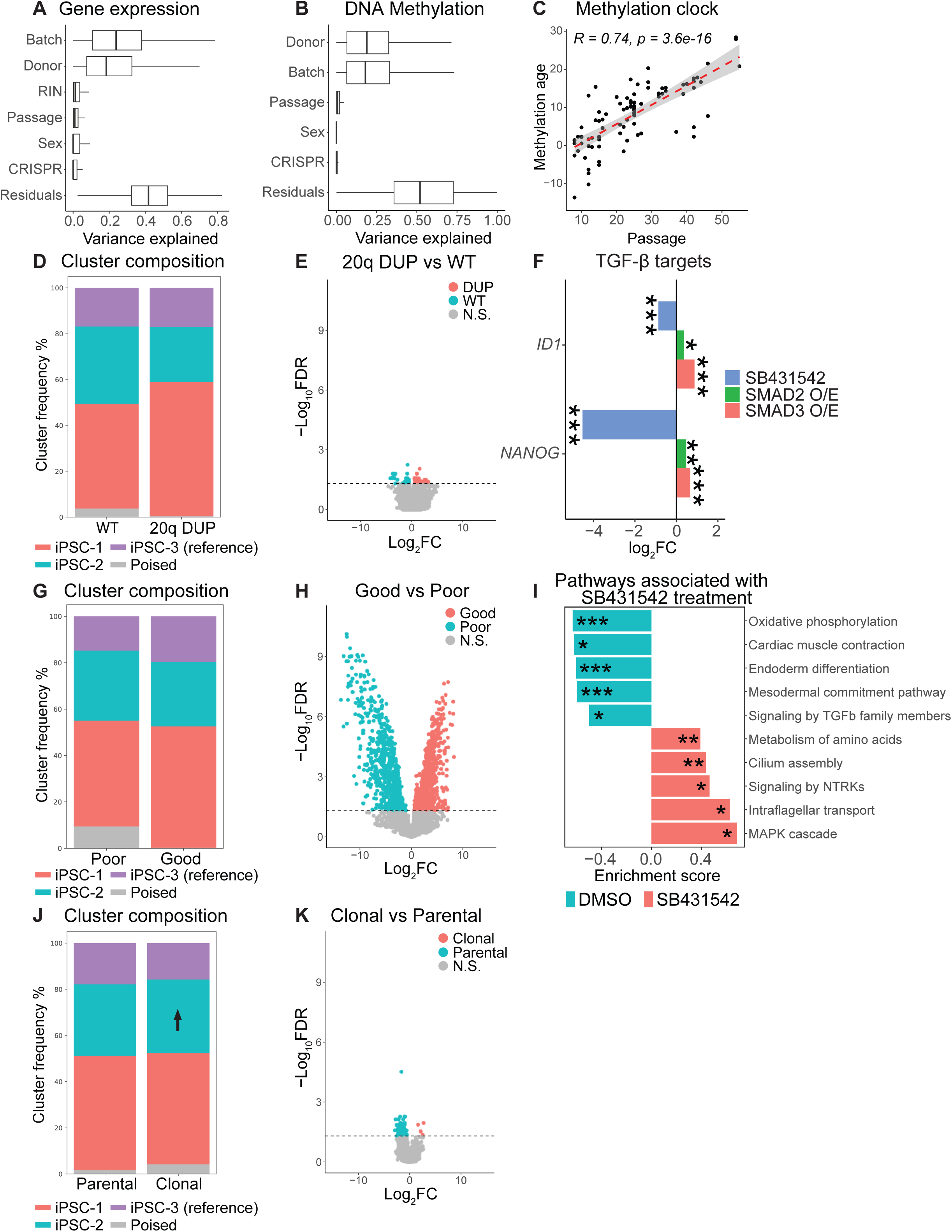
Related to Figures 4, 5 and 6: Data associated with bulk RNA-seq, cluster composition of scRNA-seq and DNA methylation array of iPSC lines (A and B) Variance partitioning of gene expression (A) and DNA methylation (B) by known technical and biological covariates using VariancePartition^56^. (C) Correlation of methylation age^66^ and the passage number of iPSC lines (D and E) Cluster composition and volcano plot of DNA methylation changes associated with 20q DUP (G) Changes in ID1 and NANOG expression after 24h SB431542 treatment^82^, SMAD2 or SMAD3 over-expression^80^ in human pluripotent stem cells. (G and H) Cluster composition and volcano plot of DNA methylation changes associated with efficient lines (I) Selected pathways associated with 24h SB431542 treatment in iPSCs^82^ using GSEA (J and K) Cluster composition and volcano plot of DNA methylation changes associated with CRISPR-editing/clonal selection See also Table S4.

